# Molecular interactome of HNRNPU reveals regulatory networks in neuronal differentiation and DNA methylation

**DOI:** 10.1101/2025.02.19.638869

**Authors:** Marika Oksanen, Francesca Mastropasqua, Krystyna Mazan-Mamczarz, Jennifer L. Martindale, Xuan Ye, Abishek Arora, Nirad Banskota, Myriam Gorospe, Kristiina Tammimies

## Abstract

HNRNPU is an RNA-binding protein with diverse roles in transcriptional and post-transcriptional regulation. Pathogenic genetic variants of HNRNPU cause a severe neurodevelopmental disorder (NDD), but the underlying molecular mechanisms are unclear. Here, we comprehensively investigate the HNRNPU molecular interactome by integrating protein-protein interaction (PPI) mapping, RNA target identification, and genome-wide DNA methylation profiling in human neuroepithelial stem cells and differentiating neural cells. We identified extensive HNRNPU-centered networks, including a strong association with the mammalian SWI/SNF chromatin-remodelling complex, and uncovered a previously unrecognized role in translation. We present evidence that HNRNPU associates with mRNAs encoding proteins important for neuronal development, including several linked to NDDs. Silencing HNRNPU reprogrammed methylation dynamics at regulatory regions, particularly at active and bivalent promoters of neurodevelopmental transcription factors. Integrative analysis across PPI, RNA, and methylome datasets identified 19 converging genes at all three molecular levels, including NDD genes within the SWI/SNF complex and RNA-processing machinery such as *SYNCRIP*. Together, these data showcase HNRNPU as a central coordinator of RNA metabolism and epigenetic remodelling during neural differentiation, linking RNA-binding, chromatin organization, and DNA methylation to the pathogenesis of HNRNPU-related NDDs.

## INTRODUCTION

Heterogeneous nuclear ribonucleoprotein U (HNRNPU), also known as scaffold attachment factor A (SAF-A), is a multifunctional protein critical for RNA metabolism and chromatin organization. HNRNPU facilitates open chromatin conformation, particularly at active promoters (1,2). Moreover, its binding near transcription start sites has been shown to increase transcriptional activity in breast cancer cell lines (3). Beyond its chromatin-related roles, HNRNPU interacts with diverse RNAs to regulate post-transcriptional processes such as RNA stabilization, splicing, and nonsense-mediated decay (4–9). In neural cells, it has been shown that HNRNPU regulates chromatin compaction, gene expression, and alternative splicing, suggesting a critical role in neural development and function (10–13). HNRNPU has also been identified in RNA-transporting granules in both human and mouse neurons (14,15) and lowering HNRNPU levels by silencing in mouse hippocampal neurons impairs RNA transport (14).

Haploinsufficiency of *HNRNPU* causes severe neurodevelopmental disorders (NDD), collectively known as HNRNPU-related NDDs, in which the affected individuals have general developmental delay, intellectual disability (ID), autism spectrum disorder (ASD), and epilepsy, among other features (16–18). Similar phenotypes have been reported in individuals having pathogenic genetic variants in other HNRNP family proteins (19). Expanding upon the genetic findings, recent epigenomic analyses have identified distinct blood DNA methylation differences in individuals with HNRNPU-related NDDs (20,21). These widespread alterations in methylation patterns suggest that chromatin dysregulation and methylation changes may underlie the neuronal progenitor survival and differentiation–processes previously implicated as central to the brain phenotypes in HNRNPU-related NDDs (10,11,13). However, methylation has not yet been investigated in neural cells.

Efforts to define the molecular interactome of HNRNPU have identified RNA targets via techniques such as ribonucleoprotein immunoprecipitation followed by RNA sequencing (RIP-Seq), formaldehyde crosslinking followed by RIP-Seq (fRIP-Seq) and enhanced cross-linking immunoprecipitation sequencing (eCLIP/CLIP-Seq) analyses in diverse cell types and tissues, including human hepatocellular carcinoma, embryonic kidney, leukemia, HeLa and pluripotent stem cell lines, as well as adrenal gland tissue (4,22–25). Additionally, two screenings to identify protein-protein interaction (PPI) partners of HNRNPU have been performed in breast cancer cell line and mouse testis tissue (3,26). Furthermore, HNRNPU has been shown to interact with multiple proteins, such as other HNRNP family members in spliceosome complexes (27). Yet, a comprehensive profiling of interacting proteins in neural cells has been lacking as previous work has focused on the effects of HNRNPU deficiency on transcriptomic changes in human neural 2D and 3D culture models (10,11) and mouse models (12,13). Here, we investigated the molecular interactome of HNRNPU during human neural development using induced pluripotent stem cell (iPSC)-derived neural cells, analyzing its PPI partners, RNA targets, and the DNA methylation landscape following HNRNPU silencing. Furthermore, we validate key interactions and their effects in an HNRNPU deficiency state, including investigation of differentially methylated regions. Through this comprehensive approach, we sought to elucidate the molecular pathways disrupted in HNRNPU-NDD models (10–13) and to understand better the mechanisms underlying the phenotypes associated with HNRNPU-related NDDs.

## MATERIAL AND METHODS

### Cell culture

The cell lines used in this study were previously established and characterized human iPSC lines derived from a neurotypical male (CTRL9-II (28), hereafter CTRL_Male_) and a neurotypical female (AF22 (29), hereafter CTRL_Female_). Both cell lines had been previously induced into neuroepithelial stem (NES) cells (29,30).

For the experiments, NES cells were seeded on 20 µg/mL poly-L-ornithine (Sigma Aldrich), and 1 µg/mL laminin (Sigma Aldrich) coated surfaces in DMEM/F12 + Glutamax medium (Gibco) with 0.05X B-27 (Gibco), 1X N-2 (Gibco), 10 ng/mL bFGF (Life Technologies), 10 ng/mL EGF (PeproTech) and 10 U/mL penicillin/streptomycin (Gibco), with daily media changes. For immunofluorescence, coverslips were coated with 100 µg/mL poly-L-ornithine and 2 µg/mL laminin. Cells were maintained in 5% CO_2_ at 37°C. NES cells were differentiated into neurogenic cultures using an undirected differentiation protocol, with media containing 0.5X B-27 and 0.4 µg/mL laminin without bFGF and EGF, with media changes every other day until day 15, then every third day. Cells were harvested at differentiation days 0 (NES, D0), 7 (D7), 14 (D14) and 28 (D28).

### Formaldehyde crosslinking and ribonucleoprotein immunoprecipitation

Formaldehyde crosslinking and ribonucleoprotein immunoprecipitation (fRIP) was performed at D0 and D28, followed by mass spectrometry (IP-MS) or RNA sequencing (fRIP-Seq) analysis. Formaldehyde crosslinking was performed as described earlier (23) with modifications. Cells were washed with DPBS, dissociated with TrypLE Express (D0) or Accutase and TrypLE Select and incubated with Neural Isolation Enzyme (D28), and resuspended in Wash medium (DMEM/F12+GlutaMAX, 5% FBS, 10 U/mL penicillin/streptomycin). Cells were gently resuspended and spun at 400 × *g* for 3 min and resuspended in Wash medium or 1 mL of PBS, supplemented with 0.15 % Triton-X and 30 μL RNase A (Sigma, R4642) to degrade all RNA as described earlier (31). Cells were counted, and wash media was added to 5×10^6^ cells/mL concentration. Cells were crosslinked with 0.1% formaldehyde for 10 min with rotation at room temperature, the reaction was stopped with 125 mM glycine for 5 min with rotation at room temperature, and washed twice in cold DPBS with 1x Halt protease inhibitor cocktail (PIC, Thermo Scientific). Cell pellets were stored at -80°C.

Cell lysis and protein immunoprecipitation were performed following an existing protocol (32) with modifications. Shortly, cell pellets were lysed in polysome lysis buffer (PLB, 100 mM KCL, 5 mM MgCl_2_, 10 mM HEPES (pH 7.0), 0.5% IGEPAL, 1 mM DTT, 100 units/mL RNaseOUT, 1xPIC). The cell lysate was incubated on ice for 10 min before centrifugation at 10,000 × *g* at 4°C. The lysate was transferred to a fresh microfuge tube and pre-cleared by incubating with Dynabeads Protein G (Life Technologies) with 25 μL of beads per sample for 30 min at 4°C with slow rotation. For fRIP-Seq, 5% of the sample was removed for input and stored at -80°C. Protein concentrations were measured using Pierce BSA protein assay kit (Thermo Scientific) and an equal protein amount of lysate was divided into two microfuge tubes per replicate, 1.5 and 1 mg for fRIP-Seq and IP-MS, respectively and NT-2 buffer (50 mM Tris (pH 7.4), 150 mM NaCl, 1 mM MgCl_2_, 0.05% IGEPAL, 1 mM DTT, 400 units/mL RNaseOUT, 16.5 mM EDTA) was added to final volume 1 mL. Either the HNRNPU antibody (ab20666, Abcam) or rabbit IgG (12-370, Sigma-Aldrich) was added at a concentration of 7 μg per 1 mg of total protein, and the reactions were incubated for 2 h at 4°C with rotation. Dynabeads protein G was prepared according to the manufacturer’s instructions, and 50 µL per 1 mg of total protein was added to fRIP-Seq and IP-MS tubes, respectively. The incubation was prolonged for 1 h. The beads were washed thrice with 1 mL ice-cold NT-2 buffer, rotating for 10 min at 4°C. After a final wash, the supernatant was removed, and beads were stored at - 80°C.

### Mass spectrometry and analysis

Mass spectrometry was performed on CTRL_Male_ samples. The beads-bound proteins were reduced in 1 mM DTT at room temperature for 30 min. Next, samples were alkylated by incubation in the dark at room temperature in 5 mM chloroacetamide for 10 min. After incubation, the remaining chloroacetamide was quenched by adding 5 mM DTT. Digestion was carried out by adding 0.4 µg Trypsin (sequencing grade modified, Pierce) and overnight incubation at 37°C. The next day, the supernatant was collected and cleaned by a modified SP3 protocol (33). Briefly, 20 µL Sera-Mag SP3 bead mix (10 µg/µL) was added to the sample. Next, 100% acetonitrile was added to achieve a final concentration of >95%. Samples were pipette-mixed, incubated for 30 min at room temperature, and then placed on a magnetic rack. The supernatant was aspirated and discarded, and the beads were washed twice in 200 µL of acetonitrile. Samples were removed from the magnetic rack, and beads were reconstituted in 20 µL phase A (97% water, 3% acetonitrile, 0.1% formic acid). Then, the beads were placed on a magnetic rack again, and the supernatant was recovered and transferred to an MS-vial. Q-Exactive Online LC-MS (LC-ESI-MS/MS) was performed using a Dionex UltiMate™ 3000 RSLCnano System coupled to a Q-Exactive mass spectrometer (Thermo Scientific); 7 µL was injected from each sample. Samples were trapped on a C18 guard desalting column (Acclaim PepMap 100, 75 µm x 2 cm, nanoViper, C18, 5 µm, 100 Å) and separated on a 50-cm long C18 column (Easy spray PepMap RSLC, C18, 2 µm, 100Å, 75 µm x 15 cm). The nano capillary solvent A was 95% water, 5% DMSO, and 0.1% formic acid; solvent B was 5% water,5% DMSO, 95% acetonitrile, and 0.1% formic acid. At a constant flow of 0.25 μl min^−1^, the curved gradient went from 6% B up to 43% B in 180 min, followed by a steep increase to 100% B in 5 min. FTMS master scans with 60,000 resolution (and mass range 300-1500 m/z) were followed by data-dependent MS/MS (30 000 resolution) on the top 5 ions using higher energy collision dissociation (HCD) at 30% normalized collision energy. Precursors were isolated with a 2 m/z window. Automatic gain control (AGC) targets were 1×10^6^ for MS1 and 1×10^5^ for MS2. Maximum injection times were 100 ms for MS1 and MS2. The entire duty cycle lasted ∼2.5 s. Dynamic exclusion was used with 60 s duration. Precursors with unassigned charge state or charge state 1 were excluded. An underfill ratio of 1% was used. The MS raw files were searched using Sequest-Percolator or Target Decoy PSM Validator under the software platform Proteome Discoverer 1.4 (Thermo Scientific) against human database and filtered to a 1% FDR cut off. We used a precursor ion mass tolerance of 15 ppm, and product ion mass tolerances of 0.02 Da for HCD-FTMS. The algorithm considered tryptic peptides with maximum 2 missed cleavage; carbamidomethylation (C) as fixed modifications; oxidation (M) as variable modifications.

For statistical analysis, only proteins identified by at least one peptide in two biological replicates were included. All keratins, histones, and immunoglobulins were considered contaminants and excluded. Significant HNRNPU-interacting proteins were selected based on FC > 2 and a Benjamini-Hochberg adjusted p-value <0.05 following a Student’s t-test. Novel interactions were identified by comparing the identified PPI list with the BioGRID database for HNRNPU interactions (accessed January 2024). Proteins were clustered using the Markov Cluster Algorithm (MCL) within the STRING database in Cytoscape (34) (v.3.10.2). Gene Ontology (GO) enrichment analysis on biological processes (BP), molecular functions (MF) and cellular components (CC) was performed using gprofiler2 package (35) (v0.2.3) in R on each cluster to identify their functional relevance, and we reported the highest significant relevant enrichment. AlphaPulldown python package (36) (v.1.0.4), which internally uses AlphaFold2 (37) was used to predict the probability of HNRNPU forming a complex with the given proteins. The input to AlphaPulldown was a compiled list of proteins pulled down with IP-MS from D0 and D28 time points. We next used the run_get_good_pae.sh command to compile a list of hits and filtered further by considering only those with interface predicted template modeling (ipTM) score > 0.6 as significant.

### RNA sequencing and analysis

fRIP-Seq was performed on samples from both CTRL_Male_ and CTRL_Female_ cell lines at D0 and from CTRL_Male_ at D28. Beads and input samples were subjected to reverse crosslinking. The beads were resuspended in 56 µL Ultra Pure water and 33 µL of reverse crosslinking buffer (PBS--, 6% N-lauroyl sarcosine, 30 mM EDTA, 7.5 mM DTT), 10 µL of Proteinase K and 1 µL of RNaseOUT were added. After incubation (1 h at 42°C then 1 h at 55°C), samples were transferred to 500 μL TRIzol for RNA extraction, including DNA digestion. An equal amount of RNA, 18 ng, was used for library preparation with Illumina Stranded Total RNA Prep with Ribo-Zero Plus, and sequencing was performed on the Illumina Nextseq 2000, P2, and 100 cycles.

RNA-seq reads were trimmed with bbduk of bbmap (v.39.01) and mapped to Homo Sapiens CRCh38 Ensembl 109 with STAR (38) (v.2.7.9a). Ribosomal reads and PCR duplicates were removed using BEDtools (v.2.31.1) intersecting with CHRCh38_rRNA.bed file (RSeQC) and Samtools (v.1.2) markdup -r, respectively. These files were subjected to CLAM (39) (v1.2.0) preprocessing, after which the peaks were called using only the uniquely mapped reads and bin size 100, read-tagger-method median, --unique-only TRUE, and –pool TRUE. Subsequent peak files were annotated with CLAM peak annotator, and resulting files were imported to R (v.4.3.2) for significance filtering. RNA with at least two peaks (adjusted p<0.05) and a BaseMean>20 in RNA-Seq data from our previous work (10) were considered significant HNRNPU targets. Enrichment of IP samples versus Input samples was analyzed using DESeq2 (40) (v.1.42.1). RNA count matrix was obtained from BAM files using Rsubread (41) (v.2.16.1). Differential expression was called using *design = ∼ Cell_Line + Status* at D0 and *design = ∼ Status* at D28, status including the information whether the sample was HNRNPU-IP or Input. A significantly enriched RNA target was called with log2FC > 1, adjusted p-value < 0.05, and base expression mean > 20. The identified RNA targets were compared with HNRNPU targets from other experiments, including fRIP-Seq performed in K562 cells (23), CLIP-Seq performed in HeLa cells (22), K562 (Encode accession ENCSR520BZQ), and HepG2 (Encode accession ENCSR240MVJ) cells, as well as adrenal gland tissue (Encode accession ENCSR368GER). The compiled list of HNRNPU targets from CLIP-Seq was downloaded from the POSTAR3 database (42). Furthermore, we compared our HNRNPU targets with our earlier RNA-seq dataset (10) (available in GEO with GSE229004), which included genes that were differentially expressed and differentially spliced in response to HNRNPU deficiency, either in a patient cell line (HNRNPUdel) or following *HNRNPU* silencing (siHNRNPU). Additional comparisons were made to external datasets, including human HNRNPU CRISPR-Cas9 mutant cortical organoids (11), and a conditional *Hnrnpu* mutant mouse model (13).

### Enrichment analyses

Expression-weighted cell-type enrichment (EWCE) analysis was performed using the EWCE package (43) (v.1.10.2) and a human neocortical mid-gestation single cell RNA-Seq (scRNA-Seq) dataset as a reference (44). The reference scRNA-Seq dataset included samples from four donors, two of which were used for this analysis. The input gene lists were either IP-MS proteins or fRIP-seq RNA targets from D0 and D28 time points separately.

Gene set enrichment analysis (GSEA) was performed using the gprofiler2 package (35) (v0.2.3) in R for gene ontology biological processes. The input gene lists were either IP-MS proteins or fRIP-seq RNA targets from D0 and D28 time points separately, and the background was set to all expressed genes (BaseMean > 20) at respective time points based on our previous RNA-seq data (10). Significant enrichment was defined as FC > 1 and adjusted p-value < 0.05. GO terms with fewer than 500 annotated genes were further filtered for semantic similarity using a *GOSemSim*-based similarity matrix and the *rrvgo::reduceSimMatrix* function with a similarity threshold of 0.6. Representative terms from the top 10 clusters were retained for visualization.

Furthermore, hypergeometric tests were performed to assess enrichment between HNRNPU PPI proteins or RNA targets and specific NDD-related gene lists: ASD (SFARI genes selected for score 1,2 and syndromic, release 08-19-2024), ID (green and amber genes from https://panelapp.genomicsengland.co.uk/ v7.0) and epilepsy (green and amber genes from https://panelapp.genomicsengland.co.uk/ v6.0). For motif enrichment analysis for RNA targets from fRIP-seq data, the adjacent CLAM peak regions were collapsed with BEDTools (v2.31.0) intersect function, and the enrichment analysis was performed with SEA function of MEME-Suite (45) (v.5.5.7) with default settings and background as shuffled input sequences. The motifs for analysis were extracted from MEME-Suite and RBPMap (v1.2) and consolidated into a list of 237 motifs from 132 proteins (Supplementary Table 1).

### Immunocytochemistry and proximity ligation assay

CTRL_Male_ cells on glass coverslips at D0 were fixed in 4% formaldehyde for 20 min at room temperature or with ice-cold methanol for 10 min. Coverslips were washed with 1xDPBS. For immunofluorescence, blocking was performed with 5% donkey serum and 0.1% Triton X-100 in 1xDPBS for 1 h at room temperature. Primary antibodies [HNRNPU (Santa Cruz Biotechnology, sc-32315, 1:25), HNRNPU (Novus Bio, NBP2-49290, 1:500), FUS (Cell Signaling Technology, 67840S, 1:1000), ARID1B (Cell Signaling Technology, 92964S, 1:1000), EEF2 (Proteintech, 20107-1-AP, 1:500), NUFIP2 (Proteintech, 67195-1-Ig, 1:1000), DDX17 (Novus Bio, NB200-352, 1:500), Fibrillarin (Cell Signaling Technology, 2639, 1:500)] were diluted in blocking buffer and incubated overnight at +4°C. After washing with 1xDPBS, secondary antibodies (Anti-Mouse 555 (Biotium), and Anti-Rabbit 488 (Invitrogen)) diluted 1:1000 and Hoechst (1:2000, Invitrogen) in blocking buffer were incubated 1 h at room temperature, protected from light. Coverslips were washed with 1xDPBS and mounted with Diamond Antifade Mountant (ThermoFisher Scientific).

The in-situ proximity ligation assay (PLA) was performed using the Duolink PLA assay kit (Sigma) with In Situ Detection Reagents Red (Sigma), following the manufacturer’s instructions. After fixation, the cells were permeabilized with 0.1% Triton in DPBS for 15 min at room temperature, blocked with supplied blocking solution for 1 h at 37°C, and incubated with primary antibodies as above. Coverslips were incubated with secondary probes (anti-mouse-PLUS and anti-rabbit-MINUS) for 1 h at 37°C, ligation solution for 30 min at 37°C and amplification solution for 100 min at 37°C, with careful washing between steps. Coverslips were mounted with Duolink in Situ Mounting Medium with DAPI. Images were acquired using LSM 900 Zeiss Confocal Microscopes using 20x or 43x magnification or LSM 980 Zeiss Confocal Microscope using 63x magnification in Airyscan mode.

### siRNA-mediated silencing

To silence *HNRNPU*, NES cells were seeded in 31,500 cells/cm^2^, left to adhere for 4 h, and transfected with 1 µM Accell SMARTpool siRNA targeting *HNRNPU* mRNA (Dharmacon E-013501-00-0010) or Accell Nontargeting siRNA (Dharmacon D-001950-01-20) according to manufacturer’s protocol. The *HNRNPU* siRNA pool consisted of the following oligos: oligo1: UCUUGAUACUUAUAAUUGU, oligo2: CUCGUAUGCUAAGAAUGGA, oligo3: GUUUCAGGUUUUGAUGCUA, oligo4: CUAGUGUGCUUGUAGUAGU. Transfected NES cells were harvested 72 h later. Cells undergoing differentiation were transfected once in the NES phase, once when switching to differentiation media, once on day 4 of differentiation, and collected on day 7 of differentiation.

### Subcellular fractionation and Western blot analysis

For total protein extraction, cells were lysed in a buffer containing 2% sodium dodecyl sulfate (SDS) in 50 mM HEPES and sonicated at 37% amplitude (Vibra-Cell VCX-600, Sonics). Subcellular fractionation was performed using the PARIS kit (Invitrogen) at differentiation days 0, 7, and 14 using CTRL_Male_ cells. Protein concentrations were measured with a Pierce BCA assay kit (Thermo Scientific). The resulting fractions were analyzed by western blot. Proteins (10 μg for total lysates and 1 μg for fractionated samples) were separated on 4-15 % gradient gels (Bio-Rad) and transferred onto nitrocellulose membranes using the Bio-Rad Turbo transfer system.

Membranes were blocked in 5% milk in TBS-Tween (0.1%) for 1 h at room temperature and incubated overnight with primary antibodies. The primary antibodies used were HNRNPU (Santa Cruz Biotechnology, sc-32315, 1:1000), Lamin B1 (Abcam, ab16048, 1:3000), GAPDH (Invitrogen, 39-8600, 1:50000), TET1 (Novus Bio, NBP2-15135 1:200) and TET3 (Proteintech, 27150-1-AP, 1:1000). After three 10-min washes in TBS-Tween at room temperature, membranes were incubated with HRP-conjugated secondary antibodies (Cell Signaling Technology: anti-mouse 7076P2, 1:2000; anti-rabbit 7074P2, 1:2000) for 1 h at room temperature. Blots were visualized using the Pierce ECL Blotting Substrate, with a 2-min incubation time, and imaged using the Bio-Rad imaging system. Western blot quantification was performed using the Fiji ImageJ software and standard densitometry techniques.

### Capillary Western blot analysis

The total protein samples (150 ng/µL) were loaded on the capillary western blot system Simple-Western-JESS (BioTechne), multiplexed for total protein and chemiluminescence detection of HNRNPU (1:50000 dilution), QSER1 (Novus Bio, NBP1-19112, 1:5 dilution), DNMT1 (Novus Bio, NB100-56519, 1:50), DNMT3A (Santa Cruz Biotechnology, sc-373905, 1:10), and DNMT3B (Santa Cruz Biotechnology, sc-376043, 1:100). Data were analyzed using Compass for S.W. software (5.0.1), and the peak areas were normalized against the total protein area. Three biological replicates were analyzed for each time point, and the Student’s t-test was used to evaluate the significance at each time point.

### RNA stability assay

To measure mRNA stability, NES cells were treated with siRNA, as described above. After 72 h of siRNA treatment, either DMSO or RNA polymerase II inhibitor actinomycin D (Sigma-Aldrich) in 5 µg/mL were added to the cells. Samples in DMSO were collected after 2 h (serving as the 0 h sample), and samples in Actinomycin D were collected in TRIzol after 2, 4, 6 or 8 h for RT-qPCR analysis. Three biological replicates were analyzed.

### Polysome profiling

Cells were incubated for 3 min with 0.1 mg of cycloheximide/mL at 37°C and then lysed in 1 mL of PEB lysis buffer (20 mM Tris-HCl [pH 7.5], 0.1 M KCl, 5 mM MgCl2, 0.5% Igepal CA-630 with RNase and protease inhibitors) as well as 0.1 mg of cycloheximide/mL to interrupt protein translation and thus prevent the disassembly of polysomes “running off” of a given mRNA. After a 10-min incubation on ice, nuclei were pelleted by centrifugation at 10,000 × *g* for 10 min at 4°C, and the supernatant was carefully layered onto a 10-to-50% linear sucrose gradient. Gradients were centrifuged at 39,000 rpm for 1.5 h at 4°C, and 1 mL fractions were collected using a system comprising a syringe pump, needle-piercing system (Brandel, Gaithersburg, MD), UV-6 detector, and fraction collector (ISCO, Lincoln, NE). The fractions were divided in half for protein samples and RNA samples (mixed with 0.5 mL TRIzol).

### RNA extraction and reverse transcription (RT) followed by quantitative (q)PCR analysis

Cells lysed in TRIzol reagent (Invitrogen) were phase-separated using chloroform and isopropanol. The aqueous phase was transferred to RNA extraction columns, and the final extraction was carried out with ReliaPrep RNA Cell Miniprep kit (Promega Z6012). The RNA was reverse transcribed using iScript cDNA Synthesis Kit (BioRad) and cDNA quantified with SsoAdvanced Universal SYBR Green Supermix (BioRad) following the manufacturer’s protocols on a CFX384 Touch Real-Time PCR Detection System (Bio-Rad). The primers used are listed in Supplementary Table 2. We used three biological replicates of cells seeded at different cell passages and reactions for each sample were performed in three technical replicates. CFX Manager software was used to record amplification curves to determine Ct values. We used three biological replicates of cells seeded at different passages. In mRNA stability assays, we calculated ΔCt relative to the levels of *GAPDH* mRNA, and ΔΔCt to samples incubated with DMSO without Actinomycin D, to determine the percentage of remaining mRNA at each time point. mRNA decay curves were modelled using polynomial regression with second-degree polynomial formula. The slope of the curve was used to determine half-life of the mRNAs. For polysome profiling, the percentage distribution for the mRNAs across the gradients was calculated using the ΔCT method.

### Click-iT protein synthesis assay

Protein synthesis after HNRNPU silencing in NES cells at D0 was measured using the Click-iT HPG Alexa Fluor 488 Protein Synthesis Assay Kit (Thermo Scientific, C10428) following the manufacturer’s instructions. Briefly, one well was pre-treated with 100 µg/mL cycloheximide for 30 min before HPG labeling to inhibit protein synthesis and confirm specificity of the Click-iT signal. All samples were labeled with 50 µM Click-iT HPG in methionine-free medium for 30 min under standard culture conditions. Cells were then fixed with 3.7% formaldehyde, permeabilized with 0.5% Triton X-100, and reacted with the Click-iT detection cocktail to visualize newly synthesized proteins. Nuclei were counterstained with HCS NuclearMask Blue Stain and coverslips were mounted and imaged using LSM 900 Zeiss Confocal Microscopes using 20x magnification, and acquisition parameters were kept constant across all samples. Quantitative image analysis was performed in ImageJ (Fiji) using a custom macro script. For each image, nuclei were segmented based on the NuclearMask Blue Stain channel using automated thresholding and watershed separation and Click-iT signal intensity was measured with the corresponding nuclear ROIs. The mean Click-iT fluorescence intensity was normalized to the number of nuclei per field of view. We used three technical replicates per condition and acquired three images per coverslip and significance was determined with the Student’s t-test.

### Whole-genome bisulfite sequencing

DNA was extracted after HNRNPU silencing using the PureLink Genomic DNA Mini kit (Thermo Scientific) following the manufacturer’s instructions. Whole-genome bisulfite sequencing (WGBS) was performed in duplicate. The DNA quality control, bisulfite conversion, library preparation, and sequencing was performed by Novogene (China). The bisulfite conversion for 100 ng input genomic DNA was performed using EZ DNA Methylation-Gold Kit (Zymo Research) and the library preparation using Accel-NGS Methyl-Seq DNA Library Kit for illumina (30096, Swift Biosciences, USA). The libraries were sequenced on NovaSeq X Plus PE150 platform, yielding on average ∼20X sequencing depth per sample. Adapter trimming and quality filtering were performed with Trimmomatic (v0.36) using the following parameters: SLIDINGWINDOW:4:15, LEADING:3, TRAILING:3, ILLUMINACLIP:adapter.fa:2:30:10, and MINLEN:36. Reads were aligned to the reference genome (Homo Sapiens CRCh38 Ensembl 109) using Bismark (46) (v0.24.0) with parameters –score_min L,0,-0.2 and -X 700 --dovetail.

Cytosine methylation levels were extracted using Bismark_methylation_extractor (--no-overlap). Differentially methylated regions were identified using DSS software with parameters smoothing.span=200, minlen = 50, minCG=3, dis.merge=100, pct.sig=0.5 and p.threshold=1×10^-5^. A significant DMR was defined as methylation difference > |0.1|. DMRs were annotated using ChIPseeker (v1.38.0), using the default priority and promoters defined as ±2000 bp. Overlaps with CpG islands, shores, shelves and open sea were annotated with CGI track from a genome cpgIslandExt and repeat regions with RepeatMasker. GSEA was performed using the gProfiler R package (35). The input consisted of either promoters overlapping DMRs or promoter DMRs overlapping H3K4me3 peaks. The universe was defined as all annotated promoters, or all promoters with H3K4me3 overlap, respectively. For visualization the terms were chosen by term size < 900 and similarity matrix was constructed as earlier described

### Cleavage Under Targets & Release Using Nuclease (CUT&RUN)

CUT&RUN was performed using CUTANA ChIC/CUT&RUN Kit v5 (Epicypher, 14-1048) following manufacturer’s protocol on extracted nuclei from untreated CTRL_Male_ cells at D0 and D7 time points. Briefly, 5×10^5^ nuclei were coupled with Concanavalin A beads with 0.01% Digitonin and incubated overnight at 4°C with 0.5 µg antibody: IgG, H3K4me3, H3K27me3 (all supplied with the kit), HNRNPU (Abcam, ab20666) or TET3 (GeneTex, GTX121453). The samples were then incubated with pAG-MNase which was activated by the addition of CaCl_2_. The reactions were incubated for 2 h at 4°C, followed by stop buffer and E. coli spike-in DNA addition. DNA was isolated using SPRI beads and sequencing libraries were prepared with 5 ng of enriched DNA using CUTANA CUT&RUN Library Prep Kit (Epicypher, 14-1001) following manufacturer’s recommendations. The libraries were sequenced on the NovaSeq X Plus PE150 platform by Novogene (Germany).

The reads were trimmed as described for fRIP-Seq reads. They were aligned to the Homo Sapiens CRCh38 Ensembl 109 using Bowtie2 (47) (v2.5.4). Up to two valid alignments were reported (-k 2) and the --dovetail option was used. Only uniquely mapped reads were retained by removing reads with secondary alignments (XS tag). Reads overlapping ENCODE DAC blacklist regions were excluded using BEDTools (v2.31.0) and duplicate reads were identified and removed as described earlier. Peak calling was performed using MACS2 (48) (v2.2.9.1) with IgG files as control. Broad peaks were identified using the parameters –broad and --broad-cutoff 0.05. To avoid model building, the –nomodel option was applied and read extension and shift were set to 300 bp and 150 bp, respectively. To identify reproducible peaks between biological replicates, peaks called from each replicate were intersected using BEDTools. Peaks were considered reproducible if they overlapped by at least 50% of their length in both replicates (-f 0.5 -r -u). The resulting consensus peak sets were used for downstream analyses. The peaks were annotated using ChIPSeeker R package (v1.38.0) using default settings.

### Statistical analysis

Statistical analyses were performed using R (v.4.3.2) and details for specific statistics are explained in the respective methods sections.

## RESULTS

### HNRNPU protein partner screen identifies six protein networks in neural cells and highlights its involvement in translation dynamics

To identify PPI partners of HNRNPU during neuronal differentiation, we performed HNRNPU IP (Supplementary Figure S1A) followed by LC-ESI-MS/MS detection of the protein lysates, with or without RNase treatment. In this study, we defined direct interactions as protein-protein contacts that remain after RNase treatment, whereas RNA-assisted interactions correspond to proteins that dissociate upon RNase digestion. Experiments were performed at two stages of differentiation: proliferative neural progenitor state at day 0 of differentiation (D0) and early differentiating neurons at day 28 (D28), which display extensive neurite outgrowth and network formation but have not yet established fully mature synaptic connections. We identified a total of 279 interacting proteins, with 244 detected at D0 and 79 at D28 (Figure 1A, Supplementary Table 3), likely reflecting the decreasing HNRNPU expression levels or function during differentiation. Among these, 72 (30.0%) and 16 (20.3%) proteins were identified as direct interactions at D0 and D28, respectively, with 13 proteins shared between both time points. When comparing our data with the BioGRID database (downloaded January 2024), we identified 54 novel direct interacting partners of HNRNPU. Analysis of the proteins dissociated after RNase treatment revealed that 172 and 63 proteins interacted with HNRNPU in an RNA-dependent manner at D0 and D28, respectively, with 31 common to both time points. Of these, 92 represented novel PPIs with HNRNPU. In total, 112 of the previously reported PPIs are RNA-assisted rather than direct (Supplementary Table 3).

**Figure 1:**
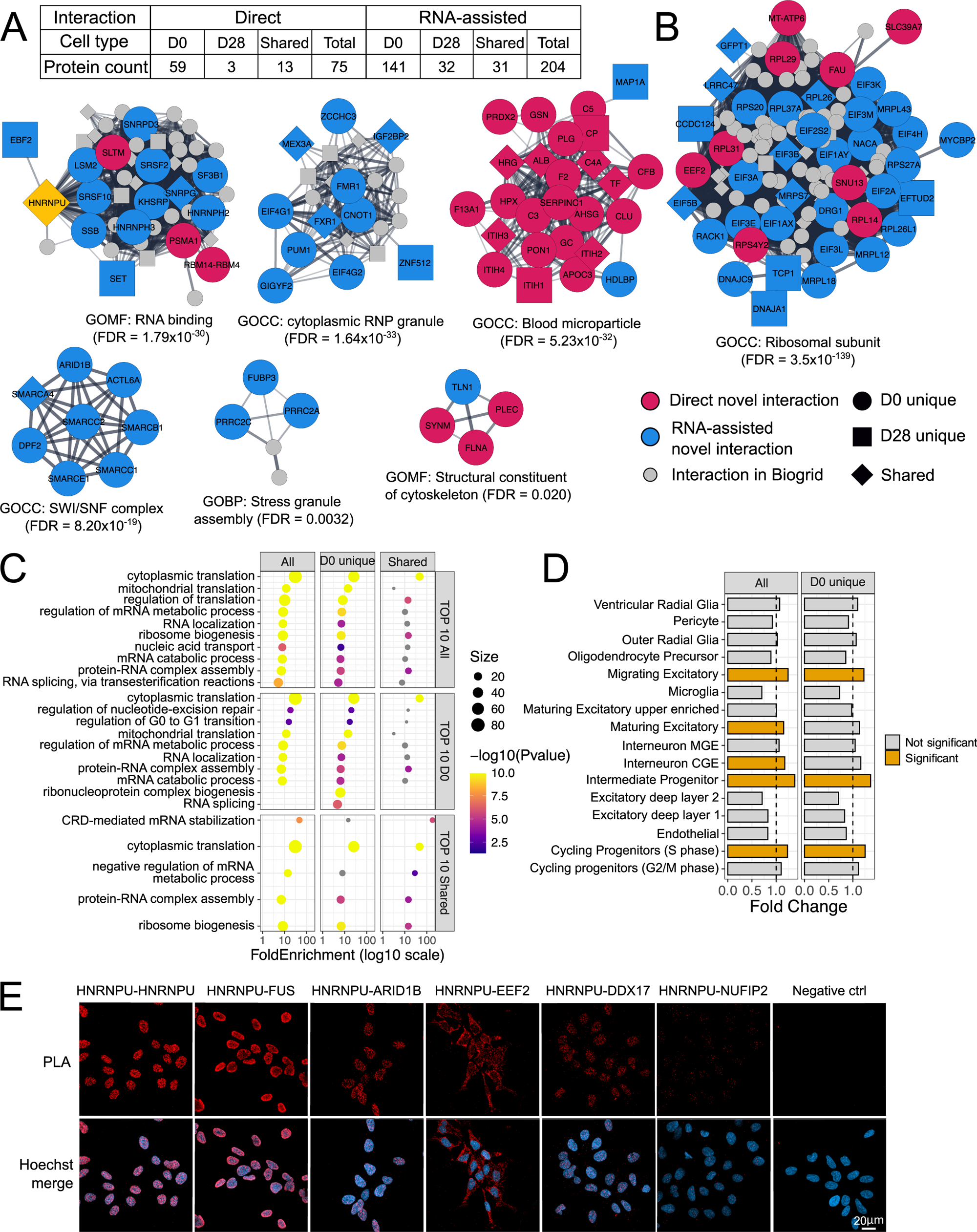
Identification and characterization of HNRNPU protein-protein interaction networks in neural cells. A) Number of HNRNPU PPI partners identified by IP-MS at D0 and D28. Proteins are classified as direct interactions (detected after RNase treatment) or RNA-assisted interactions (interaction dissociated after RNase treatment). For each category, counts are shown for proteins unique to D0, unique to D28, or shared between both time points. B) Interactome map of HNRNPU-interacting proteins based on STRING-derived protein-protein interactions. Node color indicates whether the interaction was novel and classified as direct (red) or RNA-assisted (blue), while grey nodes represent proteins with previously reported interactions in BioGRID. Only networks with significant GO enrichment are shown; all other PPIs are listed in Supplementary Table 3. For each displayed network, the most significantly enriched and relevant GO term is indicated. C) GO biological process enrichment analysis from all HNRNPU interacting proteins at D0, after merging direct and RNA-assisted interactions. D) Cell type enrichment analysis of all PPI partners identified at D0 into fetal brain mid-gestation scRNA-Seq data. E) Proximity Ligation Assay (PLA) panel representing HNRNPU interaction with the selected interacting partners. HNRNPU-HNRNPU samples were incubated with two different anti-HNRNPU antibodies. Negative control represents cells incubated without primary antibodies. Scale bar, 20 μm.

As a complimentary analysis, we used Alphapulldown to evaluate all 279 identified PPI partners. Of these, 32 interactions (11.5%) were predicted *in silico* to be direct, including HNRNPU multimerization, which has been described earlier (1) (Supplementary Table 4). The most significant proteins predicted to interact directly with HNRNPU were DEAD-box helicases DDX17 and DDX5, which we also identified at D0 (Supplementary Table 3).

Among the identified interacting partners, MCL clustering in Cytoscape revealed several protein clusters. We performed GO enrichment analysis on these clusters and identified seven protein network hubs with significant enrichment: Ribosomal subunit (FDR=3.5×10^-139^), RNA binding (FDR=1.8×10^-30^), Cytoplasmic RNP granule (FDR=1.64×10^-33^), Stress granule assembly (FDR=0.0032), Structural constituent of cytoskeleton (FDR=0.020) and SWI-SNF complex (FDR=8.20×10^-19^) (Figure 1B). Interestingly, none of the mammalian SWI-SNF (mSWI/SNF, also known as BAF) (mSWI/SNF, also known as BAF) complex proteins were earlier indicated as interacting partners of HNRNPU in the BioGRID database.

The GSEA of GO terms for all HNRNPU-interacting proteins was in line with identified networks, although it revealed distinct functional patterns depending on their temporal distribution (Figure 1C, Supplementary Table 5). Cytoplasmic translation was among the most significant terms for the combined list of all PPIs, as well as for D0-unique and shared PPIs, whereas only a single term reached significance for D28-unique proteins (negative regulation of translation, FDR=0.0049). mRNA metabolic process was among the top terms for all PPIs and D0-unique PPIs, while negative regulation of mRNA metabolic process was the most significant term for shared proteins. Furthermore, to place the identified HNRNPU PPIs in a developmental context, we performed EWCE analysis using a mid-gestation human brain scRNA-seq as a reference dataset (44). This approach assesses whether the genes encoding the identified proteins are preferentially expressed in specific cell types in the reference dataset. All identified proteins and D0-unique proteins were significantly enriched in progenitor populations with high HNRNPU expression, including cycling progenitors (S phase), intermediate progenitors, and migrating excitatory neurons, with additional enrichment in maturing excitatory neurons and CGE-derived interneurons for the full list of PPIs (Figure 1D, Supplementary Figure S1C, Supplementary Table 6). In contrast, no significant enrichment was observed for shared or D28-unique proteins. Finally, only the full set of HNRNPU PPIs was significantly enriched in ASD genes. No significant enrichments were observed for ID or epilepsy gene lists for any of the protein subsets, despite many of the proteins being encoded by causative NDD genes such as the mSWI-SNF complex (49), PUM1 (50) and ATAD3 (51) (Supplementary Figure S1D, Supplementary Table 7).

We validated four direct PPIs (FUS, EEF2, NUFIP2, and DDX17) and one RNA-assisted PPI (ARID1B) at D0 with PLA (Figure 1E). Proteins were selected for validation based on biological relevance and technical considerations: FUS served as a positive control due to its previously reported interaction with HNRNPU, EEF2 was chosen as a regulator of translation and ARID1B to represent the SWI/SNF complex. DDX17 was selected for its functional similarity to DDX5, a known HNRNPU partner, whereas NUFIP2 served as a weaker interactor with a low peptide count in our IP-MS dataset. We found that HNRNPU-EEF2 interaction occurs in the cytoplasm despite the predominant nuclear localization of the HNRNPU protein (Figure 1E). In contrast, HNRNPU-HNRNPU signal was restricted to the nucleus, which we attribute to the fact that one antibody predominantly recognizes nuclear HNRNPU, while the other also detects the cytoplasmic pool. We confirmed the presence of HNRNPU in the cytoplasm using super-resolution microscopy (Supplementary Figure S1D) and found that the cytoplasmic HNRNPU fraction increases at D7 and decreases by D14, whereas the nuclear fraction decreases in concordance with the whole fraction and our earlier RNA-Seq data (10) (Supplementary Figure S1E, Supplementary Figure S5).

### HNRNPU binds mRNAs encoding proteins related to RNA processing and neuronal development

Next, we sought to identify RNA targets of HNRNPU using fRIP-Seq at D0 and D28. We identified 21054 and 10567 significant peaks using CLAM peak calling software (39), annotated to 1707 and 734 individual genes for D0 and D28, respectively (Supplementary Table 8). Of these, 395 genes shared hits across both time points. The majority of the targets were protein-coding mRNAs (95.0% and 88.5% for D0 and D28, respectively) and long non-coding RNAs (3.5% and 5.7%), whereas the fraction of snoRNA increased from D0 (0.6%) to D28 (3.1%) (Figure 2A). The HNRNPU RNA-binding motif (UGUAUUG) was detected in 92.8% and 91.6% of the mRNA peak region sequences at D0 and D28, respectively, although the enrichment was not significant (FDR 0.849 and 0.806 at D0 and D28, respectively) (Supplementary Table 9). We compared our data to published fRIP-Seq (23) and CLIP-Seq datasets (42) from non-neural cell types and identified 1.9% and 34.4% overlap between target genes, respectively, with the latter overlap being significantly higher than expected by chance (Fisher’s exact test, p-value=2.2×10^-16^)(Figure 2B).

**Figure 2:**
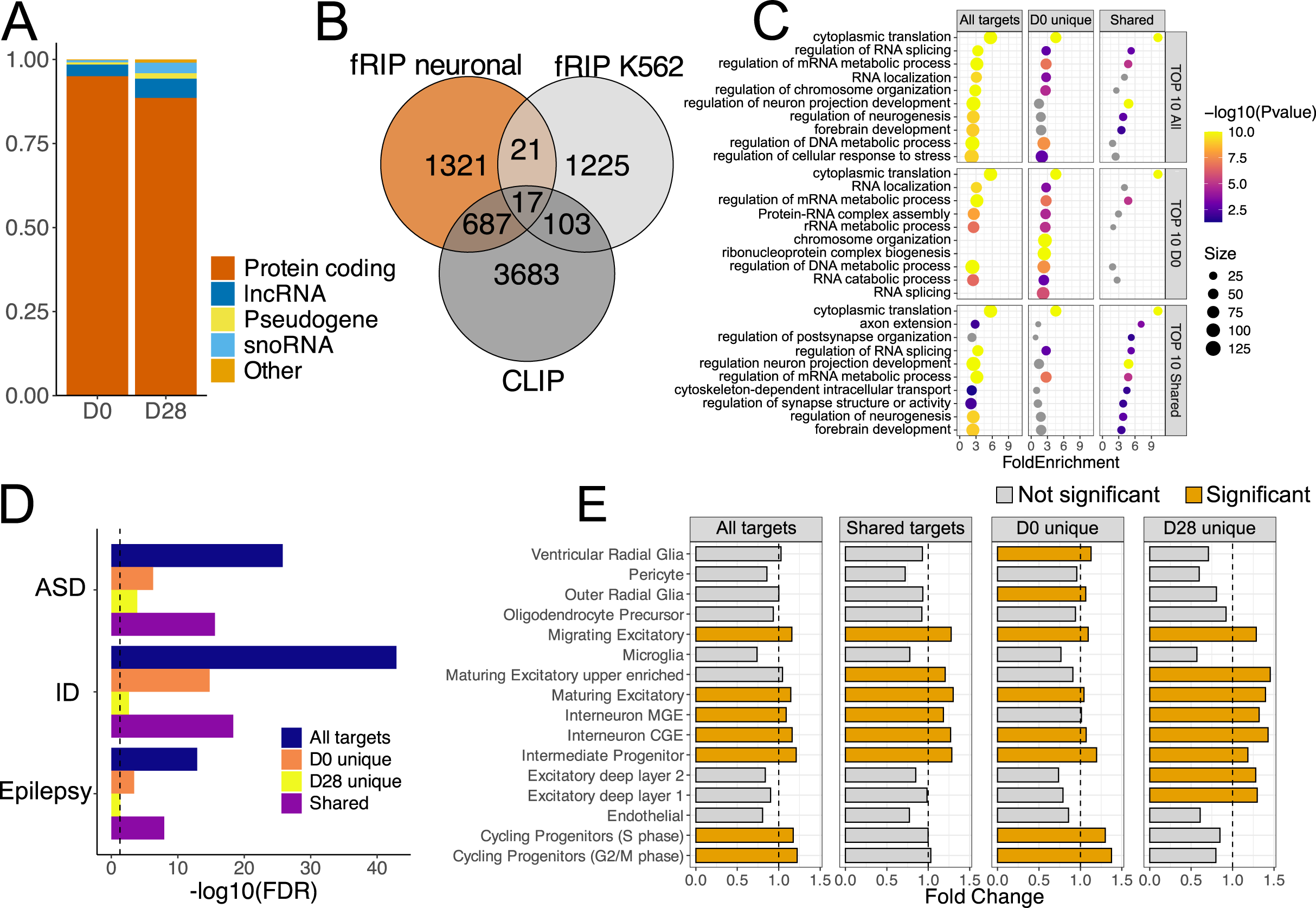
HNRNPU RNA targets and their enrichment in neurodevelopmental processes. A) Fraction of HNRNPU target RNA biotypes at D0 and D28. Biotype “Other” represents miRNA, snRNA and misc_RNA. B) Overlap of HNRNPU targets from our fRIP-Seq datasets at D0 and D28 (fRIP neuronal) and published fRIP-Seq dataset on K562 cells (23) (fRIP K562) and eCLIP/HITS-CLIP datasets from HeLa, K562, and HepG2, as well as human adrenal gland tissue (CLIP). C) GO biological process enrichment analysis on all HNRNPU target RNAs combined for D0 and D28, unique targets at D0 and shared targets at D0 and D28. Represented are the top 10 significant terms from each analysed set. D) Enrichment of the host genes of HNRNPU target RNAs combined, unique and shared at D0 and D28 in autism (ASD), intellectual disability (ID) and epilepsy risk gene lists. The vertical dotted line represents the significance threshold of FDR<0.05 (-log10(FDR)>1.3) after hypergeometric analysis. E) Cell type enrichment analysis of HNRNPU target RNAs combined, unique and shared at D0 and D28 into fetal mid-gestation scRNA-Seq data.

The full list of HNRNPU targets at D0 and D28 revealed GO-BP enrichment in cytoplasmic translation (FDR=2.04×10^-92^). The RNA targets unique to D0 time point showed additional enrichment for RNA splicing (FDR=1.24×10^-6^) and ribonucleoprotein complex biogenesis (FDR=8.61×10^-11^) among other terms, while at D28 the unique targets were only enriched in axon development (FDR=0.04) and regulation of neuron projection development (FDR=0.046) with term size<500 (Figure 2C, Supplementary Table 5). Furthermore, we found significant enrichments between HNRNPU-bound RNAs and gene lists associated with ASD, ID and epilepsy (hypergeometric test, Figure 2D, Supplementary Table 7), suggesting that mRNAs bound by HNRNPU encode proteins with crucial functions for neuronal development. Similar to the PPI, we evaluated whether HNRNPU-bound transcripts are preferentially expressed in particular cell types in the mid-gestation human brain reference dataset using EWCE analysis. While shared enrichments for HNRNPU targets were identified at D0 and D28, we also found specific enrichments at each timepoint (Figure 2E, Supplementary Table 6). For instance, D0-unique targets were enriched in progenitors and ventricular and outer radial glia, while D28-unique targets were enriched in deep-layer excitatory neurons.

To identify RNA targets that could be shared targets of the HNRNPU PPI partners, we compiled a list of RNA motifs, including motifs for 132 RBPs (Supplementary Table 1), out of which 31 proteins were also pulled down in our MS-IP experiment. We identified 29 and 2 proteins having significant motif enrichment in our fRIP-seq data at D0 and D28, respectively (Supplementary Table 10). Of these, nine proteins at D0 (EIF4G2, SRSF10, HNRNPH2, HNRNPK, IGF2BP3, MSI1, EWSR1, FUS, and FUBP3) were pulled down together with HNRNPU in our IP-MS. Seven of these were identified as RNA-assisted PPIs, and two (EWSR1 and FUS) were direct HNRNPU-interacting proteins (Supplementary Table 3).

Next, we assessed whether HNRNPU targets are differentially abundant (DA) or spliced (DS) in the HNRNPU deficiency state by comparing them with available RNA-Seq data from HNRNPU-deficient human 2D and 3D models and mouse models reported previously (10,11,13). Overall, the overlap between HNRNPU binding and simultaneous DA or DS was low, which is in line with earlier work (4,9) (Supplementary Figure S2A, Supplementary Table 8). Twenty-seven targets were DA on all datasets, including *NTRK2*, *DCX*, *NEUROD1,* and *ROBO3* mRNAs. Furthermore, comparing only to our 2D model RNA-seq at D0, D5, and D28 DA mRNAs, we found several neurodevelopmentally relevant targets to be simultaneously bound by HNRNPU and downregulated in relation to HNRNPU deficiency, such as *PAX3* and *PLXNA2* mRNAs at D0, as well as *RORB*, *NEUROD1,* and *ROBO3* mRNAs at D28. On the contrary, *LIN28A* and *PRTG* mRNAs at D0 and *CDK6*, *SOX11* and *NES* mRNAs at D28 were upregulated and bound by HNRNPU. Out of 314 DA mRNAs, 62 and 56 were also DA in *HNRNPU*-mutant 3D and mouse models, respectively (Supplementary Table 8). Similarly, in our 2D model, several bound and DS targets at D28 included mRNAs encoding proteins with critical neurodevelopmental functions, including *ANK2*, *ANK3*, *DCLK1*, *GPC1* and *NCAM1* mRNAs. Furthermore, out of 153 bound and DS targets, 31 were also DS in the mouse model, such as *NRXN1* mRNA (Supplementary Table 8).

### Functional impacts of HNRNPU on mRNA stability and translation of selected targets

Next, we aimed to investigate the effect of HNRNPU binding to those mRNAs highly enriched in the fRIP-Seq data (Figure 3A, Supplementary Table 11) that encode proteins with known relevance to neuronal development and were neither differentially spliced nor differentially abundant in HNRNPU deficiency in our published RNA-Seq data, namely *TRIM2*, *KLF12*, *ERBB4, CDK6* and *TNPO1* mRNAs. We also included *QSER1* mRNA, which has no known connection to neuronal development but was the second most significant target of HNRNPU and was recently shown to protect bivalent promoters from hypermethylation by cooperating with TET1 (52). We confirmed that silencing HNRNPU did not affect the levels of *TRIM2*, *KLF12*, or *ERBB4* mRNAs, but significantly increased the levels of *TNPO1* mRNA, and non-significantly increased *QSER1* and *CDK6* mRNAs (Figure 3B, Supplementary Figure S3A). HNRNPU silencing stabilized *TRIM2*, *QSER1* and *ERBB4* mRNA, slightly modified *KLF12* mRNA stability and had no effect on the stabilities of *TNPO1* or *CDK6* mRNAs nor that of *NPM1* mRNA, which is not an HNRNPU target (Figure 3C, Supplementary S3B). We also tested the effect of HNRNPU depletion on QSER1 protein levels and observed a nonsignificant upregulation at D0 (p-value=0.52) and D7 (p-value=0.16) (Figure 3D).

**Figure 3:**
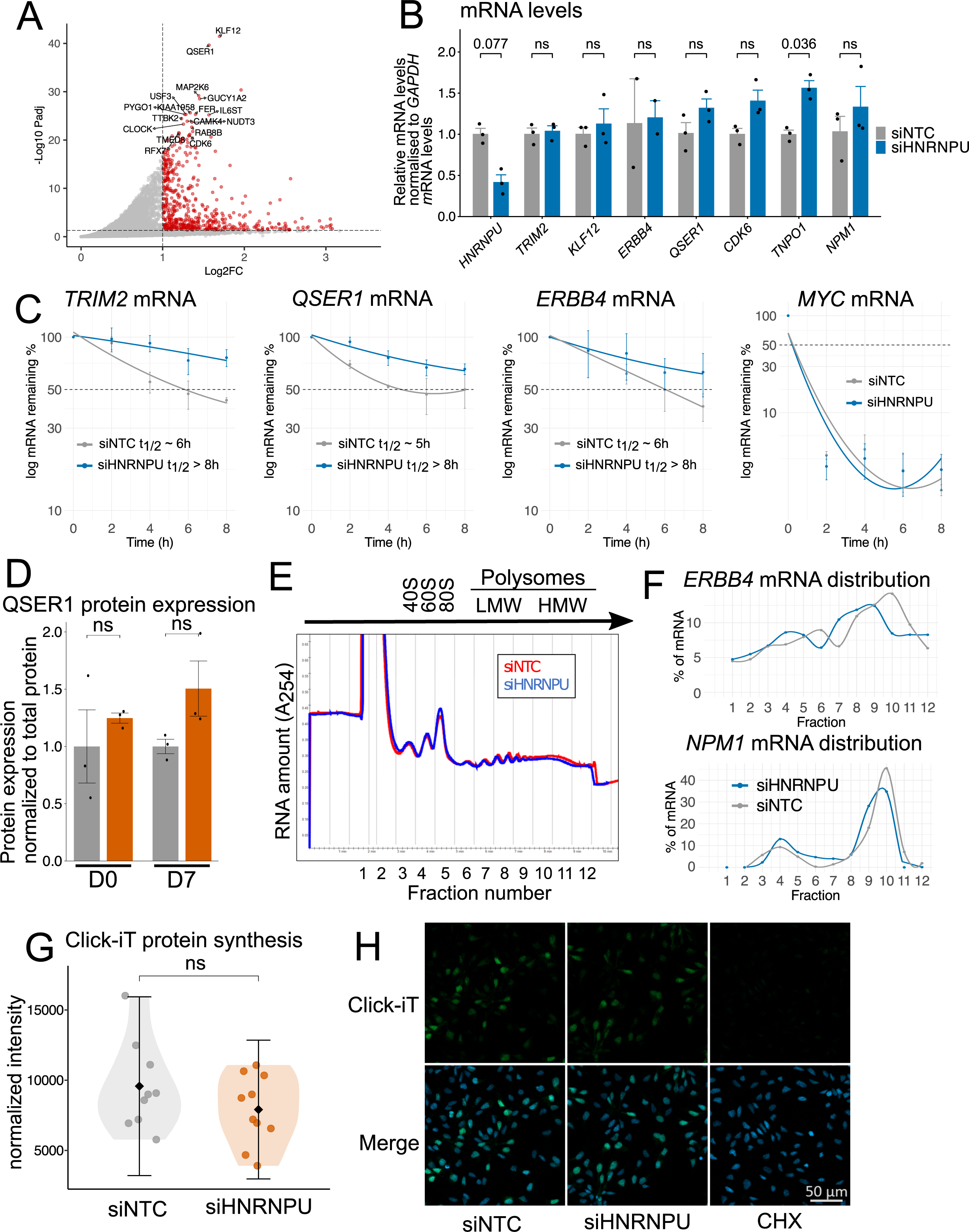
Impact of HNRNPU silencing on mRNA stability and translation in neural cells. A) A volcano plot representing DESeq2 enrichment analysis on fRIP-Seq data to identify the most abundantly enriched RNA targets of HNRNPU at D0. B) RT-qPCR analysis of the levels of *HNRNPU* mRNA, the HNRNPU targets *TRIM2*, *KLF12*, *ERBB4*, *QSER1, CDK6* and *TNPO1* mRNAs, as well as a non-target *NPM1* mRNA at D0 after HNRNPU silencing (n=2 or 3). Expression levels were normalized to the levels of *GAPDH* mRNA. Error bars represent standard error of the mean (SEM). C) RT-qPCR analysis of the stability of HNRNPU targets *TRIM2*, *QSER1* and *ERBB4* mRNA as well as a labile control *MYC* mRNA at D0 cells after HNRNPU silencing and actinomycin D treatment for the indicated times. The levels of target mRNAs were normalized to the levels of *GAPDH* mRNA. Error bars represent SEM. D) Quantification of QSER1 protein expression after HNRNPU downregulation, measured by JESS capillary western blot analysis, using total protein for normalization. Represented are fold changes relative to siNTC samples. Significance was calculated using Student’s t-test (n=3). E) Cytoplasmic lysates after HNRNPU downregulation were fractionated through sucrose gradients to evaluate polysome profiles. Fractions 1-2 represent fractions without ribosomal material, 3-5 correspond to 40S, 60S, 80S monosomes, 7-9 correspond to low-molecular-weight polysomes (LMW) and 10-12 to high-molecular-weight polysomes (HMW). F) The relative distribution of HNRNPU target *ERBB4* mRNA and a non-target *NPM1* mRNA across the fractionated polysome gradients was calculated by RT-qPCR analysis (n=1). G) Quantification of Click-iT protein synthesis assay (H). (n=9 or 10). H) Click-iT labelling of active protein synthesis. Translation inhibitor cycloheximide (CHX) was used as a control reaction. Scale bar, 50 μm.

Given the validated cytoplasmic localization of HNRNPU and the enrichment of its PPI partners and target mRNAs in translation-related processes, we next investigated whether HNRNPU regulates translation of mRNA targets. To this end, we fractionated polysomes and studied whether the relative size of polysomes forming on target mRNAs differed after modulating HNRNPU levels. The overall distribution of polysomes did not appear to vary significantly except for a slight increase in the 80S peak when HNRNPU was downregulated (Figure 3E). Polysome profiling followed by western blot analysis revealed the presence of HNRNPU in fractions one to four (corresponding to soluble fractions), and in the 40S (small ribosomal subunit), 60S (large ribosomal subunit), and 80S (monosome) fractions (Supplementary Figure S3C). Since the overall abundance of *TNPO1*, *CDK6,* and *QSER1* mRNAs increased, they were excluded from RT-qPCR analysis in polysomes.

Interestingly, the distribution of *ERBB4* mRNA across the sucrose gradients (Figure 3F) showed a shift away from the higher translating polysomes (fractions 10-12) toward lower-molecular-weight polysomes (fractions 7-9), indicating reduced translation engagement of this transcript and suggesting a potential role for HNRNPU in regulating the translation of specific mRNA targets. Furthermore, we observed slightly elevated percentages of mRNA in fraction 4 (80S peak) and fractions 7-9 (low molecular weight polysomes) for *TRIM2* and *KLF12* mRNAs in siHNRNPU samples (Supplementary Figure S3D). However, the distribution curve of non-target *NPM1* mRNA shared these same attributes (Figure 3F), suggesting these slight changes for *TRIM2* and *KLF12* mRNAs may be due to experimental conditions and do not reflect HNRNPU-specific changes.

To assess whether HNRNPU has broader effects on global translation, we employed complementary approaches. Co-staining HNRNPU and the nucleolar marker fibrillarin show that HNRNPU is absent from nucleoli, key sites for ribosomal biogenesis and assembly (Supplementary Figure S3E). Silencing HNRNPU did not affect 45S rRNA transcription or 18S rRNA maturation (Supplementary Figure S3F). Moreover, Click-iT labelling of nascent proteins revealed only a non-significant slight decrease in overall translation rates in HNRNPU-silenced cells compared with control cells (Figure 3G-H). Taken together, these results indicate that HNRNPU does not seem to regulate ribosomal biogenesis or global translation, although translation of specific target mRNAs, such as *ERBB4*, may be affected by reduced HNRNPU protein levels.

### HNRNPU deficiency reprograms promoter methylation dynamics during early neural differentiation

Given that recent studies have reported a distinct methylation signature in blood samples from individuals with HNRNPU-related NDDs (20,21), and our results demonstrate that *QSER1*, *TET1* and *TET3* mRNAs are RNA targets of HNRNPU (Supplementary Figure S4A), we next analyzed genome-wide DNA methylation patterns after HNRNPU silencing in our cell model using WGBS at D0 and D7 of differentiation (Supplementary Figure 4C). We did not observe significant differences in global methylation levels or in the distribution of methylation across genomic features between conditions (Figure 4A-B). The average CpG methylation profile within ±2 kb region around the gene bodies revealed no differences at D0 but showed hypomethylation in HNRNPU-silenced cells both upstream and downstream of gene bodies at D7 compared with control cells (Figure 4C). An overall increase in DNA methylation from D0 to D7 was expected, reflecting the establishment of the neuronal gene expression program. The relative hypomethylation observed in HNRNPU-silenced cells at D7 suggests an immature or undifferentiated epigenetic state, consistent with the delayed commitment to differentiation previously reported in HNRNPU-deficient cells (10). To assess whether altered expression of DNA methylation regulators could explain these changes, we measured protein levels of DNMT and TET enzymes following HNRNPU silencing. Although two TET3 isoforms (260 kDa and 110 kDa) appeared upregulated in siHNRNPU cells at D7, overall protein expression was highly variable and did not reach statistical significance after multiple testing correction (Supplementary Figure 4D-E, Supplementary Figure S5B-H). These findings suggest that TET3 may play a role in the observed DNA methylation alterations. However, changes in chromatin conformation following HNRNPU silencing may also contribute to these effects.

**Figure 4:**
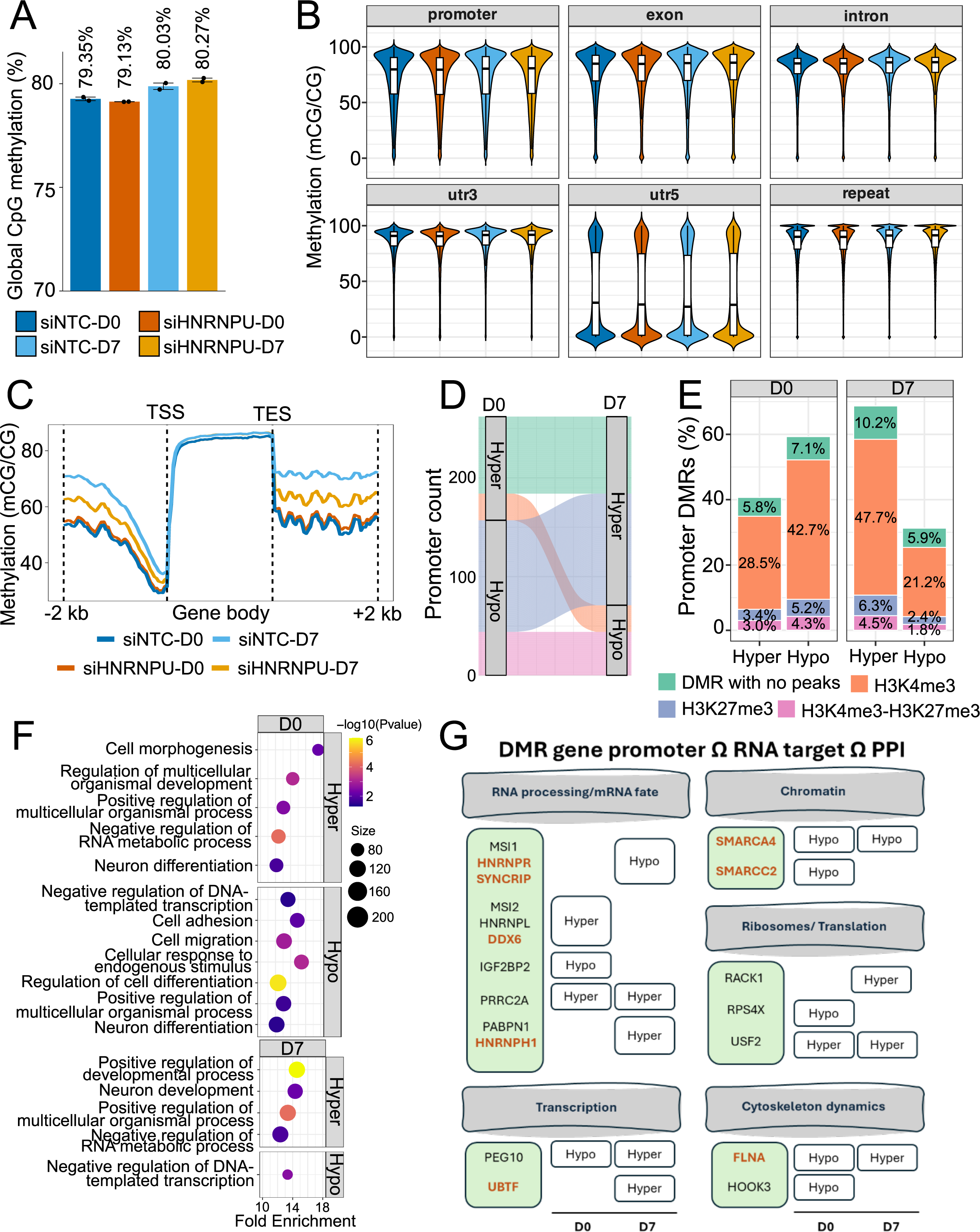
Role of HNRNPU in regulating DNA methylation and epigenetic dynamics in neural differentiation. A). Genome-wide CpG methylation levels in HNRNPU-silenced and control cells at D0 and D7 of differentiation, determined by WGBS (n=2 and n=2, respectively). B) Methylation level of genome-wide CpG methylation across different genomic features. C) Average CpG methylation profiles across gene bodies, including 2 kb upstream of the transcription start site (TSS) and 2 kb downstream of the transcription end site (TES). D) Dynamics of DMRs at promoters between D0 and D7, showing promoters that remain hypo-or hypermethylated, or switch between hypo- and hypermethylation. E) Proportion of differentially methylated promoter regions overlapping with histone modifications (H3K4me3, H3K27me3 or both) or lacking histone peaks, separated by hypermethylated and hypomethylated promoters at D0 and D7. F) GO biological process enrichment analysis on hypermethylated and hypomethylated promoter DMRs that overlapped H3K4me3 modification based on CUT&RUN analysis at D0 and D7 separately. G) Visualization of 19 genes that had differential methylation at promoters that overlapped DMRs, and the transcribed RNA is a target of HNRNPU, and the encoded protein interacts with HNRNPU protein. The genes are separated by their biological significance and highlighted in orange are genes that are known NDD risk genes. Genes in ID, epilepsy and ASD risk lists: *SMARCC2*, *HNRNPR*, *SYNCRIP*; ID and ASD: *SMARCA4*; ID: *DDX6*, *UBTF*, *HNRNPH1*, *FLNA*.

Analysis of differentially methylated regions (DMRs) revealed 3022 and 2744 DMRs at D0 and D7, respectively, out of which 1993 and 1831 overlapped gene promoters (Supplementary Table 12). Interestingly, despite the global trend toward hypomethylation around gene bodies upon HNRNPU silencing at D7, the majority of promoter DMRs were hypermethylated at this time point (68.7%), consistent with the predominant hypermethylation observed in HNRNPU-NDD patient epigenome studies (20,21). Most DMRs were unique to a single time point, but a subset of 111 promoter-associated DMRs exhibited dynamic switching, shifting from hypomethylation at D0 to hypermethylation at D7 (Figure 4D). Several promoters remained consistently hypo- or hypermethylated across differentiation. Moreover, at D0 and D7, 112 and 88 RNAs encoded by promoter-associated DMR genes were DA, with 66 and 46 showing concordant changes with methylation when compared to our earlier RNA-Seq data at D28 after HNRNPU silencing (Supplementary Table 12).

Comparing our differentially methylated CpGs (DMCs) with those shown in Rooney et al. HNRNPU-related NDD differentially methylated sites from blood samples (20) yielded no direct DMC overlaps; only one differentially methylated probe (cg16054907; FDR=0.085; Δβ=10.66%) was within 100 bp of three D0 DMCs. Similarly, no direct overlaps were found in TET3 patients comparison (53). Furthermore, we compared our promoter DMRs to promoter DMRs from Tet3-silenced mouse neural progenitor cells (54) and found 97 overlaps at D0 (60 hypo, 37 hyper; 3.27% and 2.02% of our D0 DMRs) and 93 at D7 (22 hypo, 71 hyper; 1.31% and 4.23% of our D7 DMRs), showing a D7 skew toward hypermethylation overlap.

To determine the chromatin context of the identified DMRs, we performed CUT&RUN in control cells at D0 and D7 using antibodies against H3K4me3 and H3K27me3. We also performed CUT&Run with HNRNPU and TET3 antibodies, which did not result in any reliable peaks. The majority of promoter DMRs overlapped with H3K4me3 (Figure 4E, Supplementary Table 12), indicating that these promoters were in an active chromatin state in control cells. Notably, 47.7% of hypermethylated promoter DMRs at D7 overlapped with H3K4me3, suggesting that promoters associated with active chromatin in control cells became hypermethylated and potentially associated with inactive chromatin following HNRNPU silencing. GSEA was performed separately for hypo- and hypermethylated promoter DMRs overlapping H3K4me3. While the methylation direction revealed distinct enrichment profiles, D0 and D7 showed consistent functional themes within each category (Supplementary Table 5). For example, hypermethylated promoters were associated with negative regulation of RNA metabolic process (D0 FDR=6.01×10^-5^, D7 FDR=0.014). Similarly, hypomethylated promoters consistently highlighted negative regulation of DNA templated transcription as a top term for both D0 (FDR=0.033) and D7 (FDR=0.0027).

We also identified bivalent promoters, marked by both H3K4me3 and H3K27me3, and found 634 and 559 such promoters at D0 and D7, respectively. Of these, 145 at D0 and 115 at D7 were differentially methylated, with 41.4% and 71.3% hypermethylated, respectively (Supplementary Table 12). The most hypermethylated bivalent promoter at D0 corresponded to *ZBTB7A* gene, encoding a transcription factor known to repress genes that regulate cell proliferation and differentiation (55,56). This promoter remained bivalent and hypermethylated at D7. Other crucial transcription factors with promoters associated with DMRs and marked as bivalent in our cell model included *ZBTB7C, KLF2*, *KLF4, PRDM16*, and *TCF7L1*. Together, these findings indicate that reduced HNRNPU levels disrupt DNA methylation dynamics, potentially impairing the epigenetic responsiveness required for timely differentiation.

To prioritize mechanistic candidates for further studies, we intersected promoter DMR genes with HNRNPU PPIs and RNA targets. We identified 401 genes with promoter DMRs whose transcripts were HNRNPU RNA targets (including *QSER1* and *TET3*), 40 genes with promoter DMRs encoding proteins that interact with HNRNPU, and 19 genes present in all three datasets (Supplementary Table 12). These genes recapitulate the functional enrichments shown earlier in the single datasets while providing a more restrictive set of multilayer candidates affected by HNRNPU deficiency (Figure 4G).

## DISCUSSION

Our study provides a comprehensive analysis of the molecular roles of HNRNPU, an important NDD gene, in NES and differentiating neural cells, highlighting its impact on processes affecting gene expression, including RNA metabolism, mRNA translation, chromatin remodelling complexes, and DNA methylation. Many of the found PPI partners and RNA targets were novel or not described in detail in earlier literature, highlighting the need for cell-type and state-specific interactome analysis (57,58). We also report, for the first time in this paradigm, global DNA methylation profiles together with promoter-level remodelling. Collectively, our methylation data indicate that HNRNPU deficiency selectively reprograms promoter methylation at H3K4me3-marked loci, as well as at bivalent promoters. We also showcase 19 genes identified across all levels of the molecular context of the HNRNPU interactome.

Earlier studies have focused on transcriptomic changes associated with HNRNPU deficiency. However, the direct targets of HNRNPU in RNA regulation and the initial effects of HNRNPU downregulation remained unexplored. Here, we showcase that the direct RNA targets of HNRNPU encode proteins important for neuronal development and are associated with ASD, ID, and epilepsy. Furthermore, we identified key targets that were directly bound by HNRNPU and also differentially abundant or spliced in human 2D and 3D cultured cell models, as well as in a mouse *in vivo* knockout model. For instance, *DCX* mRNA, encoding a protein crucial for cortical neuronal migration (59,60), was a common differentially expressed HNRNPU target across brain development models, and its dysregulation could relate to severe cortex development defects seen in *Hnrnpu* knockout mice (13). Similarly, we identified *ROBO3* and *NEUROD1* mRNAs as HNRNPU targets with consistently dysregulated abundance across all HNRNPU deficiency models, further underscoring the critical role of HNRNPU in neuronal development. *ROBO3* mRNA encodes a member of the ROBO receptor family that has been shown to play a key role in cortical interneuron development (61). EWCE analyses of both PPIs and mRNA targets supported this notion by showing that the HNRNPU interactome includes factors normally expressed in more advanced neural cell types *in vivo*. This suggests that HNRNPU engages early with factors that will later become critical in neuronal maturation, pointing to developmental roles that extend beyond the immediate cellular context of our model.

Given the high enrichment of translation-related proteins in our IP-MS, the verified cytoplasmic localization, and recent evidence of HNRNPU regulating translation of target mRNAs in relation to cancer (62), we investigated the effects of HNRNPU downregulation for neural target mRNAs. We found that while global translation seemed to be unaffected, HNRNPU can regulate the translation of specific mRNAs, similar to what has been shown to other RBPs such as LIN28A (63) and HNRNPC (64). One such target was *ERBB4* mRNA, encoding the tyrosine-protein kinase ERBB4 that plays a role in the central nervous system and heart development and is also associated with NDDs (65). It was noteworthy that the processes related to translation were significant both for the PPI partners and RNA targets of HNRNPU. This observation could indicate a robust regulatory loop that controls the translation machinery and protein synthesis levels of specific targets. Moreover, the downregulation of ribosomal proteins was reported as one of the most prominent biological processes downregulated in HNRNPU mutant human cortical organoids (11). To date, there have not been any proteomic-wide studies done in HNRNPU models, so further studies should provide a more comprehensive evaluation of the effects using high-throughput methods such as translatome analyses (66).

Finally, our findings show that HNRNPU deficiency alters DNA methylation during neural differentiation, specifically at promoters with an active chromatin state, supporting its role in chromatin regulation. Mechanistically, further work is needed to elucidate if there is a connection between chromatin remodelling and the observed methylation changes. However, it is noteworthy that these effects could, at least in part, arise from the strong PPI network of HNRNPU with the mSWI/SNF complex, followed by regulation of *TET3* and *QSER1*, both differentially methylated at their promoters and RNA targets of HNRNPU. The striking similarity in episignatures and clinical phenotypes between HNRNPU-related NDDs and syndromes caused by mutations in mSWI/SNF complex genes (Coffin-Siris syndrome (49)) and *TET3* (Beck-Fahrner syndrome (67)) further strengthen our hypothesis.

Specific members of the mSWI/SNF complex, *SMARCC1*, *SMARCC2* and *SMARC4A,* appear in multiple parts of the HNRNPU interactome. Notable, *SMARCC2* and *SMARCA4* promoters are also differentially methylated in relation to HNRNPU silencing. These interactions and epigenetic priming may contribute to the multifaceted role of HNRNPU in controlling key transitions during neural development. For instance, previous studies have shown that *SMARCC1* (BAF155) is involved in the transition from neural progenitors to differentiated neurons, with interacting transcription factors like PAX6, to regulate cortical development and progenitor cell division (68). On the other hand, *SMARCC2* (BAF170) is shown to be needed for the maturation of neural cells (69). SMARCA4 (BRG1), the catalytic ATPase subunit of the BAF complex, is essential for establishing chromatin accessibility at neuroectoderm enhancers and for “opening” the NPC chromatin landscape during early specification. Depletion of *BRG1* dorsalizes neural progenitor cells and promotes precocious neural crest differentiation, demonstrating its central role in lineage specification (70). Considering earlier evidence of HNRNPU localizing near transcription start sites and active promoters (1–3), we hypothesize that HNRNPU, together with the mSWI/SNF complex, facilitates access for transcription factors, DNMTs, and TET enzymes to specific DNA regions dynamically throughout early neuronal development to ensure correct transitions and differentiation patterns (71,72). Moreover, a recent study also detected interactions between HNRNPU and mSWI/SNF proteins in pluripotent stem cells (25), suggesting this interplay may play roles beyond neuronal development. Further studies are needed to test these hypotheses. Finally, we identified a set of developmentally critical TFs with promoters that showed DMRs after HNRNPU silencing, which were marked as bivalent by CUT&RUN analysis. Such promoters are typically “poised” for rapid activation or repression in response to differentiation cues, and altered methylation could compromise this flexibility. For instance, the *PRDM16* promoter was hypomethylated, whereas the *POU4F1* promoter was hypermethylated after HNRNPU silencing at D7. PRDM16 is known to promote the transition from early to late neurogenesis (73), and its altered methylation could disrupt the timing of progenitor progression. POU4F1, on the other hand, is required for the repression of early neurogenic regulators (74), and its dysregulation may impair the switch toward mature neuronal identities. In addition, we identified a subset of 19 genes showing overlap across promoter methylation, RNA binding and PPI layers, representing potential key effectors of HNRNPU loss in neural development. These multilayer candidates may form the regulatory axis through which HNRNPU coordinates chromatin remodeling, transcriptional regulation and RNA metabolism during neural differentiation. Notably, IGF2BP2, one of the genes in this subset, has been identified as a hub gene in ASD-related gene networks, with the IGF2BP1-3 complex proposed to regulate transcriptional programs associated with ASD (75).

Despite having found many novel roles and strengthened the earlier described roles of HNRNPU, we acknowledge several limitations. The HNRNPU protein is highly expressed in neural stem cells at D0, making it challenging to silence it efficiently. A challenge that is not only seen in oligo-based silencing approaches as the expression has been shown to be variable even with genome-engineered loss-of-function mutations (11). We also acknowledge that the 2D cell model used in this study is not a comprehensive representation of human brain development, and more work is needed to understand the dynamic regulation of HNRNPU in more complex 3D brain development models.

In conclusion, we present a comprehensive mapping of the HNRNPU molecular interactome and methylation changes in neural cells. Our study highlights the critical role of HNRNPU in neuronal development through its interactions with RNA regulatory pathways, including regulating translation, chromatin remodeling complexes, and DNA methylation regulators. Further studies should focus on the mechanistic understanding of the effects of the interaction modules on the observed methylation and transcriptomics changes as well as provide information at the proteomics level to fully understand the role of HNRNPU in neural development and underlying pathogenesis of *HNRNPU*-related NDDs.

## DATA AVAILABILITY

All processed data underlying this article are available as described below, except for raw sequencing data that contains potentially identifiable human data. The IP-MS data have been deposited to the ProteomeXchange Consortium via the PRIDE partner repository with the dataset identifier PXD061718. The anonymized fRIP-Seq, CUR&RUN, and WGBS data have been deposited in ENA under accession ERP170082 and ArrayExpress under accession E-MTAB-14910.

## FUNDING

This work was supported by grants from the Swedish Research Council [2017-01660 to K.T., 2023-02111 to K.T.]; Swedish Foundation for Strategic Research [FFL18-0104 to K.T.]; the Swedish Brain Foundation – Hjärnfonden [to K.T.].; KI-NIH PhD program for PhD students [to M.O.]; Osk. Huttunen Foundation [to M.O.]; the H.K.H. Kronprinsessan Lovisas förening för barnasjukvård och Stiftelsen Axel Tielmans minnesfond [2020-00573 to F.M., 2021-00617 to F.M., 2023-00784 to M.O., 2024-032 to M.O.]; the Strategic Research Area Neuroscience (StratNeuro) at Karolinska Institutet [to K.T.]; KI Foundations [to K.T.]; the Committee for Research at Karolinska Institutet [to K.T.]., Funding for open access charge: Karolinska Institute: Karolinska Institutet. K.M.M., J.L.M., N.B., and M.G. were supported by the NIA IRP, NIH.

## CONFLICT OF INTEREST

The authors have no conflicts to declare.

## ACKNOWLEDGEMENTS

The data handling was enabled by resources in project NAISS provided by the National Academic Infrastructure for Supercomputing in Sweden (NAISS) at UPPMAX, funded by the Swedish Research Council through grant agreement no. 2022-06725. We thank the PDC Center for High Performance Computing, KTH Royal Institute of Technology, Sweden, for providing access to the computing resources used in this research. We also would like to thank the core facility at NEO, BEA, Bioinformatics and Expression Analysis, which is supported by the board of research at the Karolinska Institute and the research committee at the Karolinska hospital. Mass spectrometry analysis was performed by the Clinical Proteomics Mass Spectrometry facility, Karolinska Institutet/ Karolinska University Hospital/ Science for Life Laboratory. The graphical abstract was created using Biorender.com.

## AUTHOR CONTRIBUTIONS

Conceptualization: M.O., F.M., K.M.M., M.G., and K.T.; Methodology and Data Analysis: M.O., F.M., K.M.M, J.M., X.Y., A.A., N.B., M.G., and K.T.; Validation: M.O., F.M., and K.T.; Resources: M.G. and K.T.; Data Curation: M.O., F.M,; Writing – Original Draft: M.O., and K.T.; Writing – Review & Editing: M.O., and K.T.; Visualization: M.O., and K.T.; Supervision: M.G., and K.T.; Project Administration: M.O., F.M., K.M.M., M.G., and K.T.; Funding Acquisition: M.O., and K.T., Critical review of the draft: All authors.

## SUPPLEMENTARY DATA

Supplementary Data are available at NAR online.

---

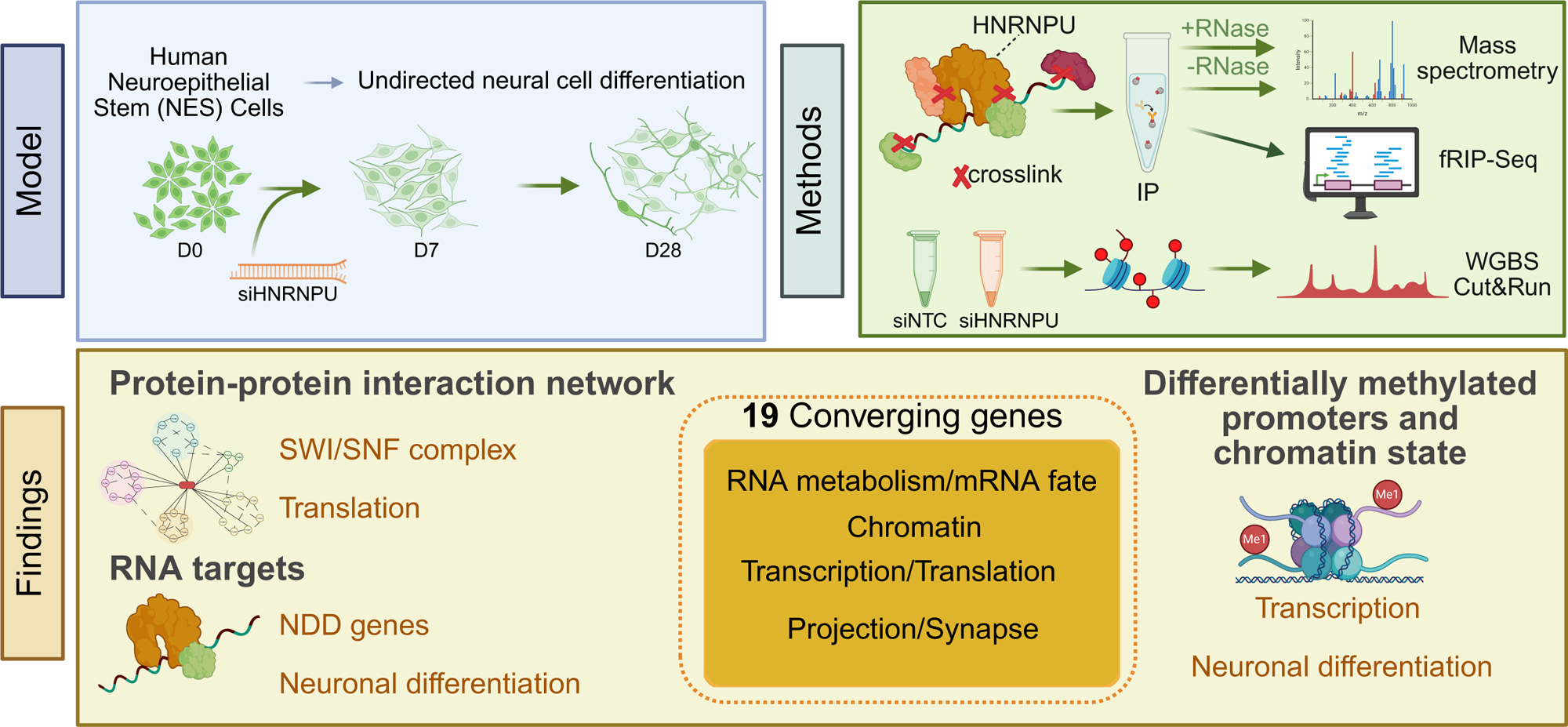

## Notes

### Competing Interest Statement

The authors have declared no competing interest.

### Summary of Updates

In revising the manuscript, we undertook substantial new analyses and data integration across all our major datasets. Specifically, we: -Added new whole-genome bisulfite sequencing (WGBS) and CUT&RUN experiments (H3K4me3 and H3K27me3) to pinpoint promoter-level methylation changes and chromatin states. -Re-analyzed and unified protein-protein interaction, RNA target, and DNA methylation datasets across datasets and across timepoints to highlight shared and unique features across differentiation stages, including a new integrated figure and expanded discussion. -Expanded explanations on cell models, validation strategies, and biological context, addressing all reviewer questions point-by-point.

