## Supplementary File 1 for "Molecular interactome of HNRNPU reveals regulatory networks in neuronal differentiation and DNA methylation"

Oksanen et al.

### Supplementary Material

#### Content

Figures S1-S5

Tables S1-S12

### Supplementary Figures

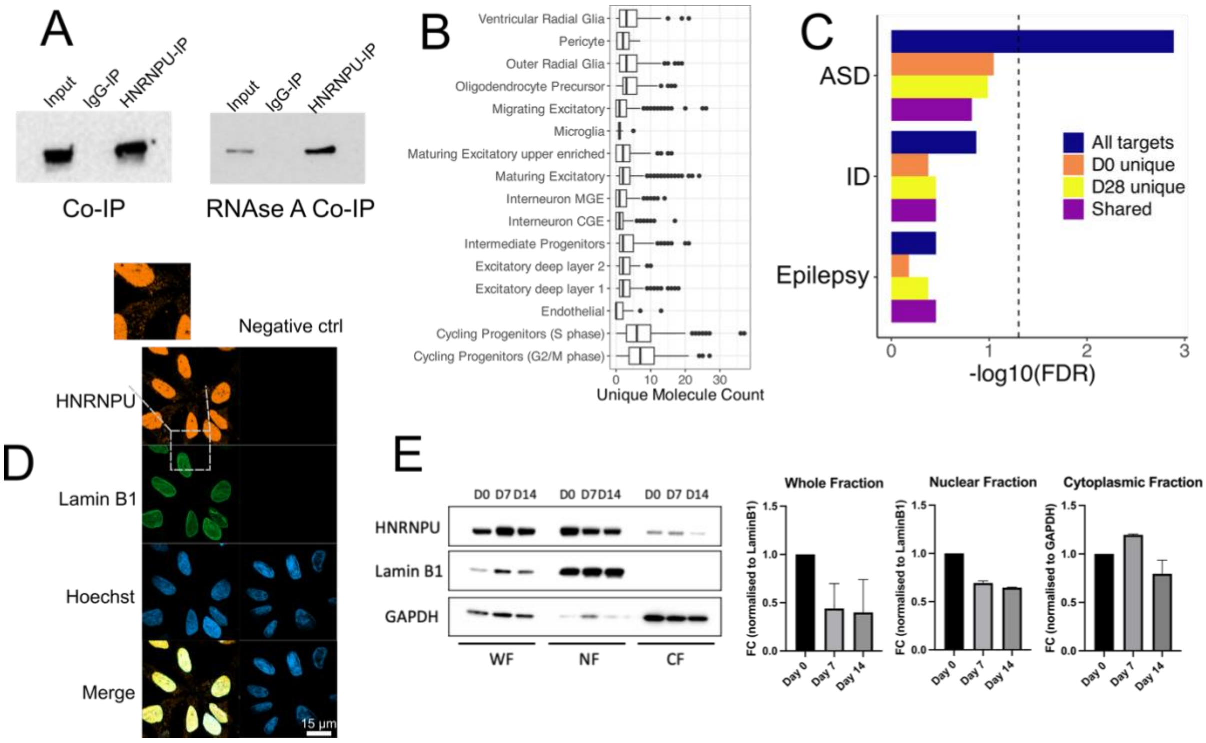


**Figure S1:** Characterization of HNRNPU protein-protein interactions and cytoplasmic localization A) Western blot analysis of untreated input sample (Input) and after Co-IP with either mock IgG control (IgG) or HNRNPU (IP). Same setting was used for Co-IP after RNase A treatment (right). B) *HNRNPU* mRNA expression per cell type in the scRNA-seq reference data set used for cell type enrichment analysis. C) Enrichment of the PPI partners of HNRNPU at D0 and D28 in autism (ASD), intellectual disability (ID) and epilepsy risk gene lists. The vertical dotted line represents the significance threshold of FDR<0.05 (-log10(FDR)>1.3) after hypergeometric analysis. D) Immunocytochemistry staining of HNRNPU, Lamin B1 and Hoechst in $CTRL_{Male}$ cells fixed with methanol, imaged with super resolution microscope. Scale bar 15 μm. E) Western blot and densitometry quantification after subcellular fractionation of $CTRL_{Male}$ cells after 0, 7 and 14 days in differentiation. WF, whole-cell fraction; NF, nuclear fraction; CF, cytoplasmic fraction. Represented are HNRNPU, nuclear marker Lamin B1 and cytoplasmic marker GAPDH. Whole-cell fraction and nuclear fraction were normalized to Lamin B1 expression and cytoplasmic fraction to GAPDH expression. (n=2)

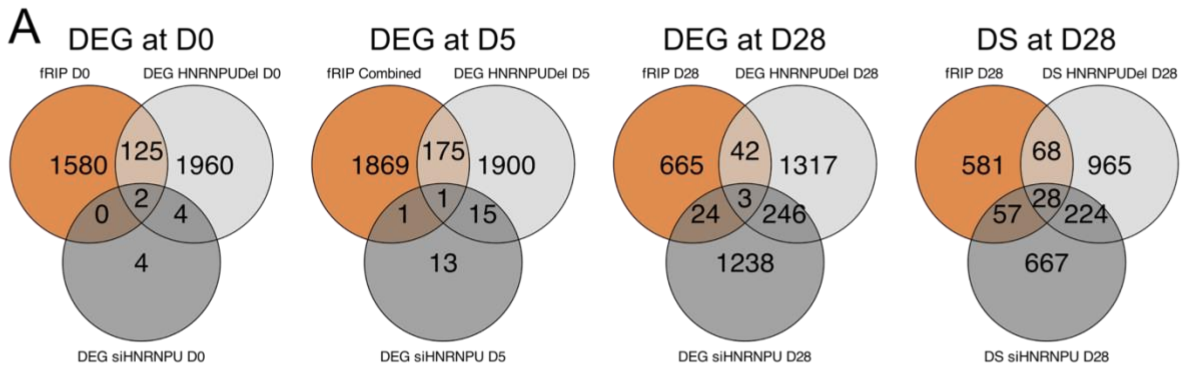


**Figure S2:** Overlap of HNRNPU RNA targets with differential expression and splicing in HNRNPU deficiency A) Venn diagrams representing overlaps between HNRNPU targets that are simultaneously differentially expressed (DE) at D0, D5 and D28, or differentially spliced (DS) at D28 in relation to *HNRNPU* deficiency. A combined list of HNRNPU targets at D0 and D28 was used when comparing with DE genes at D5, as fRIP-Seq was prepared only at D0 and D28. RNA-Seq data were from our earlier publication (Mastropasqua et al., 2023). siHNRNPU refers to siRNA silenced samples and HNRNPUdel refers to a patient cell line with heterozygous deletion of *HNRNPU*.

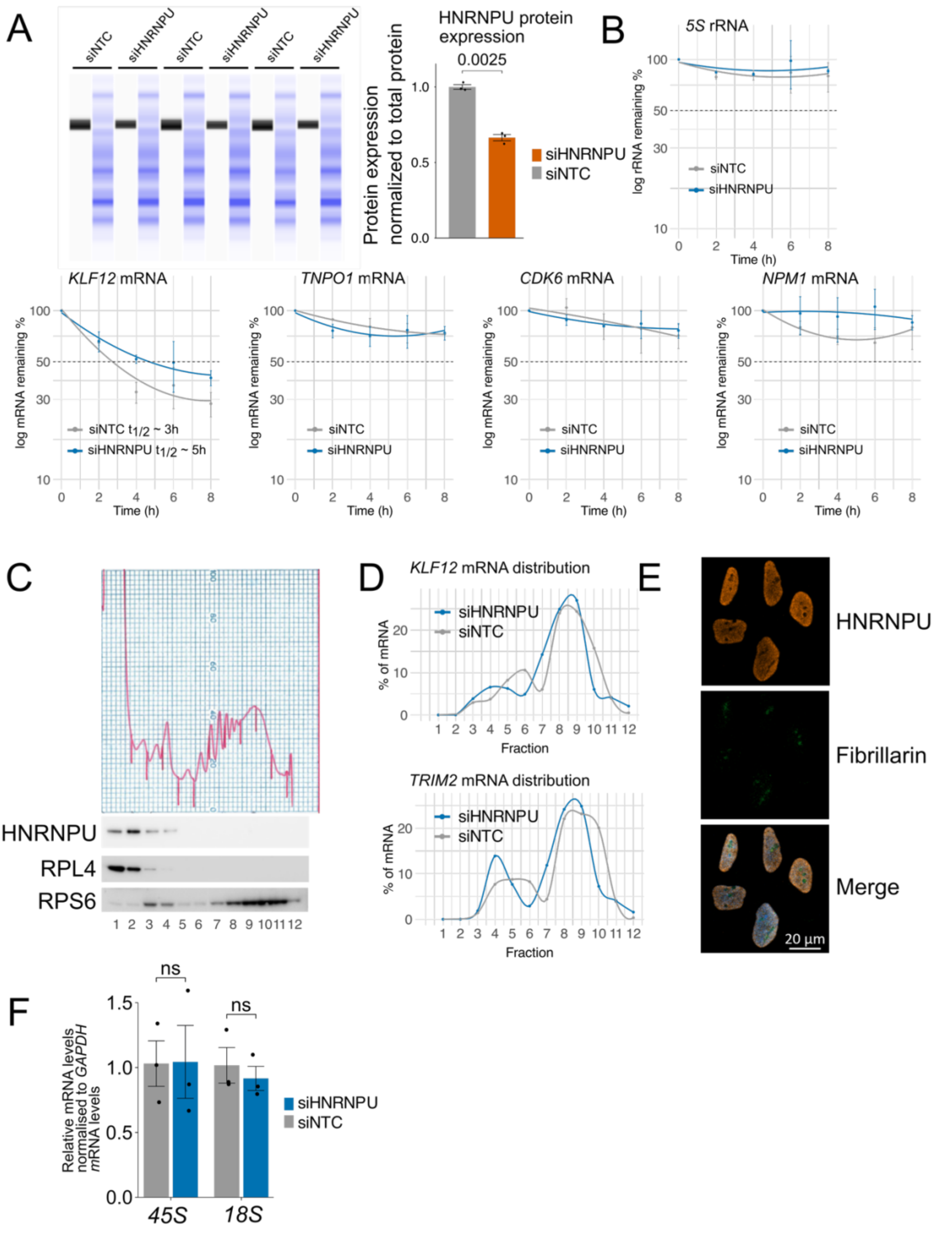


**Figure S3:** Impact of HNRNPU knockdown on mRNA stability and translation in neural cells. A) Capillary western blot and quantification of HNRNPU expression after siRNA silencing at D0. Data were normalized to total protein and represented as fold change relative to siNTC samples. Error bars represent SEM and statistical significance was calculated using Student's T-test followed by Benjamini-Hochberg multiple correction (n=3). B) RT-qPCR analysis of the stability of a stable control *5S* rRNA, HNRNPU targets *KLF12*, *TNPO1* and *CDK6* mRNAs as well as a non-target *NPM1* mRNA at D0 after HNRNPU silencing and actinomycin D treatment for the indicated times (n=3). C) Polysome profiling at D0, followed by HNRNPU, RPL4 and RPS6 western blot analysis. Fractions 1-2 represent fractions without ribosomal material, 3-5 are 40S, 60S, 80S monosomes, 6-8 low-molecular-weight polysomes and 9-12 high-molecular-weight polysomes. D) Relative mRNA distribution of HNRNPU targets *KLF12* and *TRIM2* mRNAs on polysome gradients as assessed by RT-qPCR analysis (n=1). E) Immunocytochemistry staining of HNRNPU, nucleoli marker Fibrillarin and Hoechst in $CTRL_{Male}$ cells imaged with a confocal microscope. Scale bar 20 μm. F) Quantification of *45S* rRNA and *18S* rRNA levels after HNRNPU silencing measured by RT-qPCR. Error bars represent SEM and statistical significance was calculated using Student's T-test followed by Benjamini-Hochberg multiple correction (n=3).

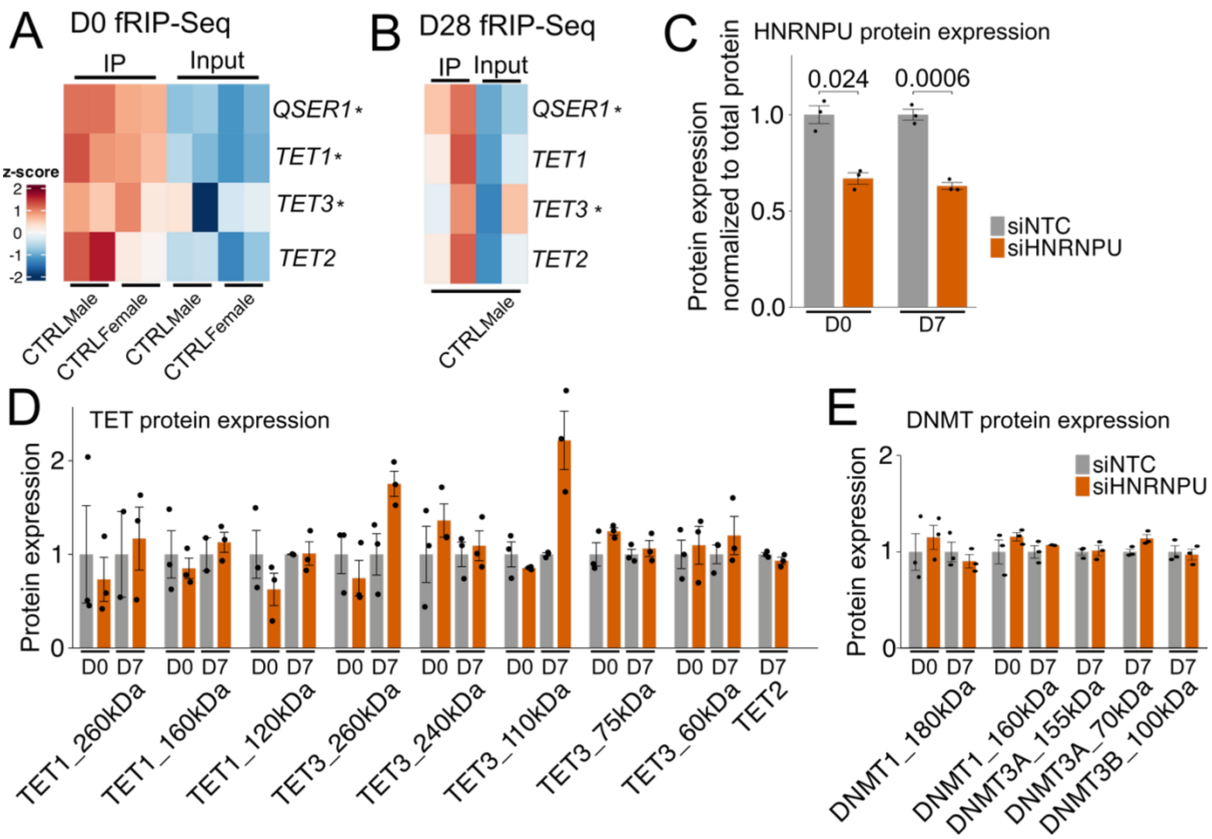


**Figure S4:** HNRNPU targets mRNAs encoding DNA methylation regulators in differentiating neural cells A-B) Heatmap of *QSER1*, *TET1*, *TET3* and *TET2* mRNAs from fRIP-Seq data from $CTRL_{Male}$ at D0 (A) and D28 (B). Asterisks represent significant HNRNPU targets. *TET3* mRNA was a significant HNRNPU target based on CLAM peak calling analysis at D28. B) Quantification of HNRNPU expression after silencing at D0 and D7 measured by capillary western blot. Data were normalized to total protein and represented as fold change relative to siNTC samples. Error bars represent SEM, and statistical significance was calculated using Student's t-test followed by Benjamini-Hochberg multiple correction (n=3). D-E) Western blot and capillary western blot quantification of TET1/2/3 (D) and DNMT1/3A/3B (E). Data were normalized to total protein for capillary western blot and either Lamin B1 or GAPDH for western blot and represented as fold change relative to siNTC samples. Western blots and images of capillary western blots are presented in Figure S5. Error bars represent SEM, and statistical significance was calculated using Student's t-test followed by Benjamini-Hochberg multiple correction (n=3).

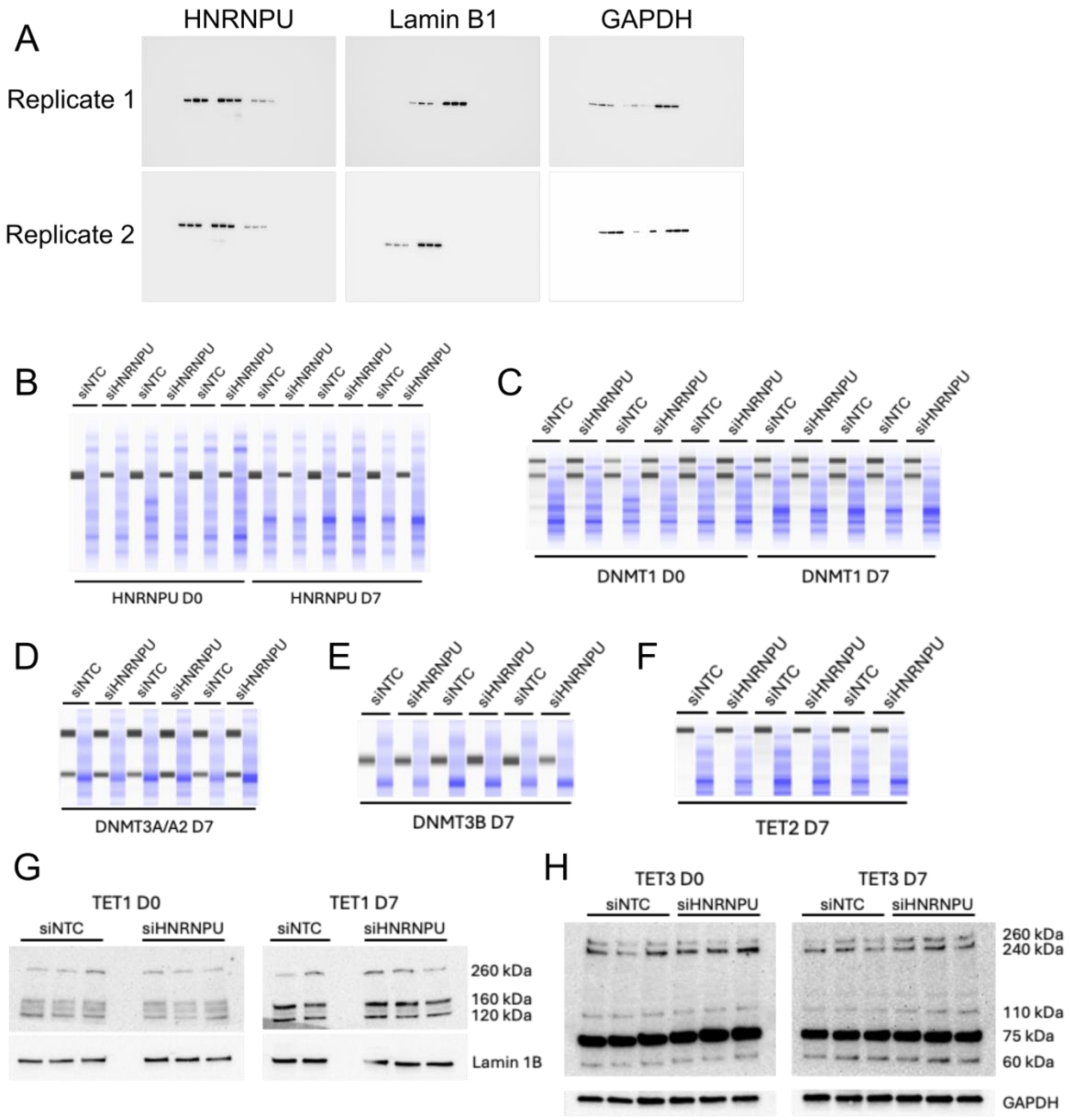


**Figure S5:** Full western blot and capillary western blot images. A) Full images of western blots related to figure S1. Represented are HNRNPU, nuclear marker Lamin B1 and cytoplasmic marker GAPDH, from two biological replicates. B-H) Capillary western blot images representing HNRNPU (B), DNMT1 (C), DNMT3A/A2 (D), DMT3B (E), and TET2 (F) either at both D0 and D7 or only D7 time points. G-F) Full images of western blots of TET1 (G) and TET3 (F) at D0 and D7 time points.

### Supplementary Tables

**Table S1.** A list of motifs compiled from MEME-Suite (v.5.5.7) and RBPMap (v.1.2), used in motif enrichment analysis of fRIP-Seq data. See separate file.

**Table S2.** Primers used in RT-qPCR analysis

| mRNA | Forward | Reverse |
|---|---|---|
| *5S* | CGCCCGATCTCGTCTGAT | GGTCTCCCATCCAAGTACTAACCA |
| *CDK6* | GGCTCTAACCTCAGTGGTCGT | CAACACTCCAGAGATCCACGG |
| *ERBB4* | TAGTTCAGGATGTGGACGTTGC | ACACACCGTCCTTGTCAAAGT |
| *GAPDH* | AAGGTGAAGGTCGGAGTCAAC | GGGGTCATTGATGGCAACAATA |
| *HNRNPU* | AGTTTAACAGAGGTGGTGGCC | GCCCCTCCTATTATATCCGCC |
| *KLF12* | CAGTCGGTGCCTGTTGTCTA | GGTCCATTTGTGCTTTGCCA |
| *NPM1* | GCTGGTGCAAAGGATGAGTTG | CCAAGGGAAACCGTTGGCT |
| *MYC* | TTCTCTCCGTCCTCGGATTCTCTG | TCTTCTTGTTCCTCCTCAGAGTCG |
| *QSER1* | CACCTCAGCCTACTGGTCTT | CCTGAGCAGTTCGGGATTCA |
| *TNPO1* | GGCCTGACCTCTTACCAAAAC | ACCAAATGCTCCCTCACAG |
| *TRIM2* | TGAAGGCACCAACATCCCAA | GTAGTTCTGCAGGCACCTCTC |
| *45S* | TCGTGCTGCCCTCTCGG | GGAACGACACACCACCGTT |
| *18S* | CGAACGTCTGCCCTATCAACTT | ACCCGTGGTCACCATGGTA |

**Table S3.** List of significantly enriched HNRNPU protein-protein interacting partners at D0 and D28. Column 'Interaction_Type' specifies whether the interaction was classified direct or RNA-assisted based on RNAse A treated samples, while value 'direct, RNA-assisted' indicates that the interaction was direct at D0 and RNA-assisted at D28. Column 'Time_Point' specifies whether the protein was interacting with HNRNPU at D0, D28 or shared between both time points. Column 'Biogrid' specifies wether the interaction with HNRNPU had been deposited to BioGrid database. Listed are peak areas, average peak areas, fold change, P values and adjusted P values for D0 and D28 samples, with and without RNAse A treatment. See separate file.

**Table S4.** Results of AlphaPulldown analysis. Significance was determined by ipTM > 0.6. Column 'HNRNPU Domain' specifies which functional domain of HNRNPU was involved in the interaction.

| HNRNPU Interactor | HNRNPU Domain | Inter-face | Num_intf residues | Polar | Hydro-phobhic | Charged | Contact pairs | sc | hb | sb | int_solv_en | int_area | pi_score | ipTM | mpDockQ /pDockQ |
|---|---|---|---|---|---|---|---|---|---|---|---|---|---|---|---|
| DDX39A | SAP | C_B | 9 | 0.111 | 0.333 | 0.444 | 8 | 0.616 | 5 | 3 | -0.91 | 461.31 | 1.04 | 0,8601 | 0,2004 |
| DDX17 | SPRY | C_B | 22 | 0.045 | 0.409 | 0.318 | 25 | 0.405 | 5 | 0 | -10.26 | 1312.03 | -0.02 | 0,8482 | 0,3517 |
| DDX5 | SPRY | C_B | 22 | 0.091 | 0.409 | 0.273 | 19 | 0.411 | 3 | 1 | -9.18 | 1123.81 | -0.36 | 0,8397 | 0,3226 |
| FBL | SAP | C_B | 11 | 0.364 | 0.273 | 0.273 | 11 | 0.578 | 3 | 4 | -0.18 | 468.91 | -0.11 | 0,8135 | 0,1860 |
| RPS4Y2 | SAP | C_B | 10 | 0.2 | 0.3 | 0.5 | 8 | 0.446 | 2 | 1 | -4.4 | 569.87 | -0.28 | 0,7940 | 0,1490 |
| EIF3L | SAP | C_B | 9 | 0.333 | 0.333 | 0.222 | 7 | 0.642 | 5 | 4 | -0.59 | 479.7 | 0.63 | 0,7911 | 0,1344 |
| EIF3C | SPRY | C_B | 17 | 0.294 | 0.588 | 0.118 | 16 | 0.501 | 1 | 0 | -7.32 | 547.05 | -0.64 | 0,7809 | 0,1271 |
| RPS4X | SAP | C_B | 10 | 0.2 | 0.3 | 0.5 | 8 | 0.435 | 3 | 0 | -4.4 | 546.81 | -0.26 | 0,7791 | 0,1420 |
| HNRNPU | ATPase | C_B | 29 | 0.448 | 0.172 | 0.241 | 29 | 0.519 | 16 | 4 | -5.25 | 1237.71 | 0.51 | 0,7646 | 0,2270 |
| GIGYF2 | ATPase | C_B | 25 | 0.32 | 0.28 | 0.24 | 24 | 0.437 | 9 | 6 | -6.6 | 1069.05 | -0.82 | 0,7614 | 0,1619 |
| SRSF2 | SAP | C_B | 9 | 0.111 | 0.222 | 0.444 | 9 | 0.504 | 5 | 2 | -0.2 | 345.37 | -0.19 | 0,7598 | 0,1467 |
| FAU | SPRY | C_B | 41 | 0.22 | 0.39 | 0.146 | 57 | 0.67 | 14 | 3 | -13.84 | 1412.91 | 2.22 | 0,7225 | 0,4704 |
| USF1 | SPRY | C_B | 27 | 0.259 | 0.481 | 0.185 | 31 | 0.582 | 8 | 0 | -14.39 | 1177.5 | 1.28 | 0,7157 | 0,1324 |
| EIF5B | ATPase | C_B | 24 | 0.125 | 0.458 | 0.25 | 19 | 0.359 | 1 | 7 | -11.23 | 1155.07 | -0.48 | 0,7149 | 0,1581 |
| SLIRP | SAP | C_B | 15 | 0.133 | 0.467 | 0.2 | 15 | 0.526 | 1 | 0 | -7.89 | 487.12 | 0.02 | 0,7148 | 0,1936 |
| RPL23A | SAP | C_B | 9 | 0.222 | 0.222 | 0.333 | 7 | 0.475 | 1 | 2 | -0.56 | 260.88 | -1.22 | 0,7120 | 0,0759 |
| SMARCB1 | ATPase | C_B | 17 | 0.353 | 0.353 | 0.118 | 12 | 0.456 | 7 | 4 | -8.68 | 962.87 | -0.68 | 0,7056 | 0,2090 |
| EFTUD2 | ATPase | C_B | 12 | 0.333 | 0.417 | 0.083 | 9 | 0.434 | 3 | 4 | -1.92 | 590.94 | -1.57 | 0,7020 | 0,0987 |
| EIF2A | SPRY | C_B | 7 | 0.143 | 0.429 | 0.143 | 4 | 0.626 | 2 | 0 | -3.17 | 712.39 | -0.53 | 0,6991 | 0,1037 |
| PDAP1 | SPRY | C_B | 20 | 0.3 | 0.25 | 0.35 | 23 | 0.425 | 6 | 4 | 1.77 | 683.2 | -1.6 | 0,6955 | 0,0876 |
| CCDC136 | ATPase | C_B | 24 | 0.292 | 0.292 | 0.25 | 18 | 0.441 | 5 | 9 | -9.57 | 1392.82 | -0.73 | 0,6856 | 0,1636 |
| SRSF10 | SAP | C_B | 11 | 0.182 | 0.182 | 0.545 | 11 | 0.613 | 3 | 2 | -0.81 | 355.87 | 0.68 | 0,6691 | 0,1443 |
| USF2 | SPRY | C_B | 46 | 0.239 | 0.413 | 0.196 | 51 | 0.443 | 12 | 0 | -24.66 | 1768.75 | 0.75 | 0,6671 | 0,1560 |
| RPL31 | ATPase | C_B | 7 | 0.143 | 0.571 | 0.286 | 5 | 0.628 | 2 | 5 | 0.74 | 341.16 | -0.2 | 0,6599 | 0,0369 |

| | | | | | | | | | | | | | | | |
|---|---|---|---|---|---|---|---|---|---|---|---|---|---|---|---|
| RIOK2 | SPRY | C_B | 87 | 0.322 | 0.356 | 0.195 | 173 | 0.076 | 22 | 1 | -17.72 | 2274.53 | 0.79 | 0,6497 | 0,5619 |
| FUBP3 | SPRY | C_B | 70 | 0.2 | 0.357 | 0.243 | 106 | 0.268 | 14 | 10 | -10.29 | 2143.99 | 0.05 | 0,6493 | 0,2939 |
| RTN1 | ATPase | C_B | 34 | 0.235 | 0.441 | 0.206 | 28 | 0.305 | 7 | 2 | -13.91 | 1331.93 | -1.27 | 0,6492 | 0,2409 |
| DAP3 | ATPase | C_B | 11 | 0.455 | 0.364 | 0.091 | 12 | 0.579 | 4 | 1 | 0.87 | 358.9 | -0.43 | 0,6436 | 0,0762 |
| SMARCC2 | SPRY | C_B | 35 | 0.257 | 0.457 | 0.171 | 39 | 0.476 | 7 | 0 | -14.87 | 1287.24 | 0.29 | 0,6434 | 0,0951 |
| LASP1 | SPRY | C_B | 28 | 0.286 | 0.464 | 0.25 | 34 | 0.19 | 5 | 5 | -17.4 | 1224.31 | -0.42 | 0,6419 | 0,0813 |
| SLTM | ATPase | C_B | 37 | 0.324 | 0.324 | 0.27 | 40 | 0.257 | 15 | 5 | -8.85 | 1395.69 | -1.11 | 0,6087 | 0,1657 |
| JPT2 | SPRY | C_B | 19 | 0.158 | 0.579 | 0.105 | 24 | 0.583 | 8 | 2 | -11.88 | 795.78 | 1.31 | 0,6007 | 0,1870 |

**Table S5.** GO biological process enrichment analysis of HNRNPU PPIs and mRNA targets at D0 and D28 and promoter DMRs that overlapped H3K4me3 CUT&RUN peaks at D0 and D7. The input list for PPIs consisted of both direct and RNA-assisted PPIs. The PPI and fRIP-Seq input lists were split by all, D0-unique, D28-unique and shared PPIs/RNA targets, while DMRs were split by all, hyper- and hypomethylated DMRs. Sheet 1: HNRNPU PPI enrichment all proteins. Sheet 2: HNRNPU PPI enrichment at D0-unique. Sheet 3: HNRNPU PPI enrichment at D28-unique. Sheet 4: HNNRPU PPI enrichment shared proteins. Sheet 5: HNRNPU RNA target enrichment all targets. Sheet 6: HNRNPU RNA target enrichment D0-unique. Sheet 7: HNRNPU RNA target enrichment D28-unique. Sheet 8: HNRNPU RNA target enrichment shared proteins. Sheet 9: All promoter DMRs overlapping with H3K4me3 enrichment at D0. Sheet 10: Hypomethylated promoter DMRs overlapping with H3K4me3 enrichment at D0. Sheet 11: Hypermethylated promoter DMRs overlapping with H3K4me3 enrichment at D0. Sheet 12: All promoter DMRs overlapping with H3K4me3 enrichment at D7. Sheet 13: Hypomethylated promoter DMRs overlapping with H3K4me3 enrichment at D7. Sheet 14: Hypermethylated promoter DMRs overlapping with H3K4me3 enrichment at D7. See separate file.

**Table S6.** Cell type enrichment analysis (EWCE) of HNRNPU PPIs (IP-MS) and RNA targets (fRIP-Seq) at D0 and D28. The input lists were separated for all proteins/targets, D0-unique, D28-unique and shared proteins/targets. See separate file.

**Table S7.** Enrichment analysis of HNRNPU PPIs (IP-MS) and HNRNPU mRNA targets (fRIP-Seq) on autism (ASD), intellectual disability (ID) and epilepsy risk gene lists (hypergeometric test). Background gene lists were obtained from our previous RNA-Seq data from Mastropasqua et al., 2023, with BaseMean > 20. Q refers to intersection of PPI and NDD risk genes, and m refers to a number of NDD risk genes. The input lists were separated for all proteins/targets, D0-unique, D28-unique and shared proteins/targets.

| Assay | Input | NDD | Background | q | m | P-value | FDR | -log10(P-value) |
|---|---|---|---|---|---|---|---|---|
| IP-MS | All proteins | SFARI | 19679 | 29 | 964 | 0,00010786 | 0,00129436 | 2,8879 |
| IP-MS | All proteins | ID | 19679 | 34 | 1785 | 0,04511947 | 0,13535841 | 0,8685 |
| IP-MS | All proteins | Epilepsy | 19679 | 13 | 749 | 0,26242334 | 0,34989779 | 0,4561 |
| IP-MS | D0 unique | SFARI | 15987 | 19 | 883 | 0,014969 | 0,089814 | 1,0467 |
| IP-MS | D0 unique | ID | 15987 | 23 | 1703 | 0,3810244 | 0,41947198 | 0,3773 |
| IP-MS | D0 unique | Epilepsy | 15987 | 8 | 705 | 0,66096173 | 0,66096173 | 0,1798 |
| IP-MS | D28 unique | SFARI | 19162 | 5 | 958 | 0,0257849 | 0,1031396 | 0,9866 |
| IP-MS | D28 unique | ID | 19162 | 5 | 1777 | 0,20333734 | 0,34989779 | 0,4561 |
| IP-MS | D28 unique | Epilepsy | 19162 | 2 | 747 | 0,38451598 | 0,41947198 | 0,3773 |
| IP-MS | Shared proteins | SFARI | 19679 | 5 | 964 | 0,06270102 | 0,15048245 | 0,8225 |
| IP-MS | Shared proteins | ID | 19679 | 6 | 1785 | 0,20550076 | 0,34989779 | 0,4561 |
| IP-MS | Shared proteins | Epilepsy | 19679 | 3 | 749 | 0,23428524 | 0,34989779 | 0,4561 |
| fRIP-Seq | All targets | SFARI | 19679 | 212 | 964 | 2,6379E-27 | 1,5827E-26 | 25,8006 |
| fRIP-Seq | All targets | ID | 19679 | 376 | 1785 | 1,0225E-44 | 1,227E-43 | 42,9112 |
| fRIP-Seq | All targets | Epilepsy | 19679 | 145 | 749 | 6,0791E-14 | 1,2158E-13 | 12,9151 |
| fRIP-Seq | D0 unique | SFARI | 15987 | 115 | 883 | 3,4952E-07 | 5,2427E-07 | 6,2804 |
| fRIP-Seq | D0 unique | ID | 15987 | 232 | 1703 | 6,7988E-16 | 1,6317E-15 | 14,7874 |
| fRIP-Seq | D0 unique | Epilepsy | 15987 | 84 | 705 | 0,00031282 | 0,00037538 | 3,4255 |

| | | | | | | | | |
|---|---|---|---|---|---|---|---|---|
| fRIP-Seq | D28 unique | SFARI | 19162 | 34 | 958 | 9,3494E-05 | 0,00012466 | 3,9043 |
| fRIP-Seq | D28 unique | ID | 19162 | 48 | 1777 | 0,00205961 | 0,00224685 | 2,6484 |
| fRIP-Seq | D28 unique | Epilepsy | 19162 | 20 | 747 | 0,04393156 | 0,04393156 | 1,3572 |
| fRIP-Seq | Shared targets | SFARI | 19679 | 63 | 964 | 8,3903E-17 | 2,5171E-16 | 15,5991 |
| fRIP-Seq | Shared targets | ID | 19679 | 96 | 1785 | 1,1236E-19 | 4,4944E-19 | 18,3473 |
| fRIP-Seq | Shared targets | Epilepsy | 19679 | 41 | 749 | 6,3646E-09 | 1,0911E-08 | 7,9621 |

**Table S8.** List of significant HNRNPU RNA targets at D0 and D28. Sheet 1: HNRNPU mRNA targets based on CLAM peak calling from fRIP-Seq data at D0 and D28. The column 'Timepoint' specifies whether the mRNA was a target at D0, D28 or shared target at both time points. Sheet 2: HNRNPU mRNA targets from fRIP-Seq data that show either differential expression or differential splicing in HNRNPU deficient 2D (Mastropasqua et al. 2023), 3D (Ressler et al. 2023) or mouse (Sapir et al. 2022) datasets. See separate file.

**Table S9.** MEME enrichment analysis of HNRNPU RNA motif in CLAM peak regions at D0 and D28.

| Time Point | CONSENSUS | TP | TP% | FP | FP% | ENR _RATIO | SCORE _THR | PVALUE | EVALUE | QVALUE |
|---|---|---|---|---|---|---|---|---|---|---|
| D0 | UGUAUUG | 9235 | 91.64 | 9320 | 92.48 | 0.991 | 1.5 | 8.06e-1 | 8.06e-1 | 8.06e-1 |
| D28 | UGUAUUG | 18486 | 92.84 | 18575 | 93.29 | 0.995 | 1.4 | 8.49e-1 | 8.49e-1 | 8.49e-1 |

**Table S10.** MEME enrichment analysis results at D0 and D28. The motifs used were from Table S1 and the input was RNA sequences from CLAM peak regions, split by All targets, D0-unique, D28-unique and shared targets. The D28-unique analysis yielded no significant results.

| Input | Motif ID | Protein | Concensus | TP | TP% | FP | FP% | Enr ratio | Score thr | P-value | E-value | Q-value |
|---|---|---|---|---|---|---|---|---|---|---|---|---|
| All targets | RNCMPT00160 | HNRNPH2 | GGGAGGG | 2091 | 8.25 | 1447 | 5.71 | 1.44 | 8.9 | 1.11e-27 | 1.42e-25 | 7.78e-26 |
| All targets | RBPMap-FUBP1-2 | FUBP1 | UDUUU | 4344 | 17.14 | 3716 | 14.67 | 1.17 | 7.4 | 1.40e-12 | 1.79e-10 | 4.90e-11 |
| All targets | RNCMPT00086 | ZC3H14 | UUUGUUU | 1894 | 7.47 | 1501 | 5.92 | 1.26 | 8.4 | 8.19e-12 | 1.05e-9 | 1.91e-10 |
| All targets | RNCMPT00090 | SRSF10 | AGAGAMA | 373 | 1.47 | 217 | 0.86 | 1.72 | 10 | 6.91e-11 | 8.84e-9 | 1.21e-9 |
| All targets | RNCMPT00041 | MSI1 | UAGUWRG | 3723 | 14.69 | 3195 | 12.61 | 1.17 | 7.6 | 1.16e-10 | 1.48e-8 | 1.62e-9 |
| All targets | RNCMPT00117 | HuR | UUUGUUU | 1459 | 5.76 | 1137 | 4.49 | 1.28 | 9.1 | 1.41e-10 | 1.81e-8 | 1.65e-9 |
| All targets | RBPMap-ELAVL4-2 | ELAVL4 | UUWUU | 6310 | 24.90 | 5633 | 22.23 | 1.12 | 6.2 | 3.06e-10 | 3.92e-8 | 3.06e-9 |
| All targets | RNCMPT00112 | HuR | UUUGUUU | 1023 | 4.04 | 764 | 3.02 | 1.34 | 9.8 | 4.87e-10 | 6.23e-8 | 4.26e-9 |
| All targets | RNCMPT00070 | SNRNP70 | RWUCAAG | 1040 | 4.10 | 783 | 3.09 | 1.33 | 10 | 9.54e-10 | 1.22e-7 | 7.41e-9 |
| All targets | RNCMPT00044 | PCBP2 | CCUYCCC | 996 | 3.93 | 762 | 3.01 | 1.31 | 9.1 | 1.31e-8 | 1.68e-6 | 9.17e-8 |
| All targets | RBPMap-BOLL-2 | BOLL | UUUDUUU | 937 | 3.70 | 713 | 2.81 | 1.31 | 7.2 | 1.92e-8 | 2.46e-6 | 1.22e-7 |
| All targets | RNCMPT00089 | SRSF10 | AGAGARR | 208 | 0.82 | 110 | 0.43 | 1.88 | 10 | 2.11e-8 | 2.70e-6 | 1.23e-7 |
| All targets | RBPMap-ELAVL4-1 | ELAVL4 | UAAUU | 2609 | 10.30 | 2240 | 8.84 | 1.16 | 9.4 | 6.21e-8 | 7.95e-6 | 3.34e-7 |
| All targets | RNCMPT00088 | SRSF10 | AGAGARR | 146 | 0.58 | 69 | 0.27 | 2.1 | 9.9 | 8.13e-8 | 1.04e-5 | 4.06e-7 |
| All targets | RNCMPT00150 | ESRP2 | UGGGGAU | 1346 | 5.31 | 1090 | 4.30 | 1.23 | 9 | 1.16e-7 | 1.49e-5 | 5.42e-7 |
| All targets | RNCMPT00177 | SFPQ | GURGUKU | 439 | 1.73 | 300 | 1.18 | 1.46 | 9.2 | 1.78e-7 | 2.28e-5 | 7.79e-7 |
| All targets | RBPMap-EIF4G2-3 | EIF4G2 | GGUYGC | 1090 | 4.30 | 865 | 3.41 | 1.26 | 10 | 1.97e-7 | 2.53e-5 | 8.12e-7 |

| | | | | | | | | | | | | |
|---|---|---|---|---|---|---|---|---|---|---|---|---|
| All targets | RBPMap-EWSR1-2 | EWSR1 | GGGDGGGG | 176 | 0.69 | 94 | 0.37 | 1.86 | 11 | 3.41e-7 | 4.37e-5 | 1.33e-6 |
| All targets | RNCMPT00187 | BRUNOL6 | UGUGDKG | 504 | 1.99 | 378 | 1.49 | 1.33 | 9.9 | 1.25e-5 | 1.59e-3 | 4.58e-5 |
| All targets | RBPMap-FUBP3-1 | FUBP3 | UAUAU | 2390 | 9.43 | 2127 | 8.39 | 1.12 | 8.5 | 4.82e-5 | 6.17e-3 | 1.69e-4 |
| All targets | RNCMPT00050 | RBM3 | GAUACGA | 3894 | 15.37 | 3561 | 14.05 | 1.09 | 7 | 6.01e-5 | 7.69e-3 | 2.00e-4 |
| All targets | RNCMPT00019 | SRSF10 | AGAGAAA | 211 | 0.83 | 140 | 0.55 | 1.5 | 10 | 8.91e-5 | 1.14e-2 | 2.83e-4 |
| All targets | RNCMPT00067 | SRSF9 | GGRWGGA | 208 | 0.82 | 138 | 0.54 | 1.5 | 11 | 9.92e-5 | 1.27e-2 | 3.02e-4 |
| All targets | RBPMap-ESRP1-1 | ESRP1 | GGGKGG | 705 | 2.78 | 576 | 2.27 | 1.22 | 9 | 1.72e-4 | 2.21e-2 | 5.02e-4 |
| All targets | RNCMPT00113 | RBM4 | GCGCGSG | 2927 | 11.55 | 2661 | 10.50 | 1.1 | 6.2 | 1.96e-4 | 2.51e-2 | 5.48e-4 |
| All targets | RNCMPT00032 | HuR | UUWUUUU | 479 | 1.89 | 378 | 1.49 | 1.27 | 10 | 3.14e-4 | 4.02e-2 | 8.44e-4 |
| All targets | RBPMap-FUS-1 | FUS | GGKGG | 806 | 3.18 | 675 | 2.66 | 1.19 | 10 | 3.62e-4 | 4.64e-2 | 9.32e-4 |
| All targets | RNCMPT00162 | LIN28A | CGGAGGR | 6781 | 26.76 | 6393 | 25.23 | 1.06 | 6.1 | 3.73e-4 | 4.78e-2 | 9.32e-4 |
| All targets | RNCMPT00154 | RBM5 | GAAGGAG | 2366 | 9.34 | 2147 | 8.47 | 1.1 | 8.2 | 5.86e-4 | 7.50e-2 | 1.41e-3 |
| All targets | RNCMPT00152 | RBMS1 | UAUAUAS | 399 | 1.57 | 313 | 1.24 | 1.27 | 11 | 7.14e-4 | 9.14e-2 | 1.66e-3 |
| All targets | RNCMPT00173 | RBMS3 | UAUAUAB | 1061 | 4.19 | 919 | 3.63 | 1.15 | 9.5 | 7.62e-4 | 9.76e-2 | 1.72e-3 |
| All targets | RNCMPT00026 | HNRNPK | CCAAMCC | 348 | 1.37 | 271 | 1.07 | 1.28 | 11 | 1.11e-3 | 1.43e-1 | 2.43e-3 |
| All targets | RNCMPT00004 | BRUNOL4 | UGUGUGU | 718 | 2.83 | 609 | 2.40 | 1.18 | 9.3 | 1.51e-3 | 1.93e-1 | 3.19e-3 |
| All targets | RNCMPT00056 | RBM8A | GCGCGCG | 1057 | 4.17 | 929 | 3.67 | 1.14 | 9.4 | 2.18e-3 | 2.79e-1 | 4.49e-3 |
| All targets | RNCMPT00013 | DAZAP1 | UAGGUAR | 69 | 0.27 | 40 | 0.16 | 1.71 | 12 | 3.52e-3 | 4.51e-1 | 7.03e-3 |
| All targets | RNCMPT00166 | BRUNOL5 | UGUGUGU | 634 | 2.50 | 541 | 2.14 | 1.17 | 10 | 3.63e-3 | 4.64e-1 | 7.04e-3 |
| All targets | RNCMPT00057 | RBMS3 | AUAUAUM | 454 | 1.79 | 376 | 1.48 | 1.21 | 11 | 3.74e-3 | 4.79e-1 | 7.08e-3 |
| All targets | RNCMPT00107 | SRSF1 | GGAGGAN | 2603 | 10.27 | 2415 | 9.53 | 1.08 | 8.8 | 4.14e-3 | 5.30e-1 | 7.48e-3 |

| | | | | | | | | | | | | |
|---|---|---|---|---|---|---|---|---|---|---|---|---|
| All targets | RNCMPT00064 | SART3 | ARAAAAA | 3557 | 14.04 | 3337 | 13.17 | 1.07 | 7.7 | 4.17e-3 | 5.34e-1 | 7.48e-3 |
| All targets | RNCMPT00155 | PABPC1 | ARAAAAA | 3556 | 14.03 | 3337 | 13.17 | 1.07 | 7.7 | 4.32e-3 | 5.53e-1 | 7.55e-3 |
| All targets | RNCMPT00049 | RBM28 | GWGUAGW | 226 | 0.89 | 175 | 0.69 | 1.29 | 11 | 6.22e-3 | 7.96e-1 | 1.06e-2 |
| All targets | RBPMap-FUBP1-1 | FUBP1 | UAUGUAU | 485 | 1.91 | 411 | 1.62 | 1.18 | 9.7 | 7.35e-3 | 9.40e-1 | 1.20e-2 |
| All targets | RNCMPT00171 | PABPC5 | AGAAAAU | 1231 | 4.86 | 1112 | 4.39 | 1.11 | 9 | 7.38e-3 | 9.45e-1 | 1.20e-2 |
| All targets | RNCMPT00052 | RBM4 | GCGCGGG | 4891 | 19.30 | 4656 | 18.37 | 1.05 | 3.7 | 8.31e-3 | 1.06e0 | 1.32e-2 |
| All targets | RBPMap-DAZ3-1 | DAZ3 | ACGUUU | 285 | 1.12 | 233 | 0.92 | 1.22 | 12 | 1.25e-2 | 1.60e0 | 1.94e-2 |
| All targets | RBPMap-FUBP3-3 | FUBP3 | UUUWU | 1164 | 4.59 | 1061 | 4.19 | 1.1 | 8 | 1.53e-2 | 1.96e0 | 2.25e-2 |
| All targets | RNCMPT00156 | CNOT4 | GACAGAN | 2386 | 9.42 | 2238 | 8.83 | 1.07 | 8.4 | 1.53e-2 | 1.96e0 | 2.25e-2 |
| All targets | RNCMPT00186 | PCBP1 | CCUWWCC | 3299 | 13.02 | 3125 | 12.33 | 1.06 | 8 | 1.54e-2 | 1.98e0 | 2.25e-2 |
| All targets | RNCMPT00109 | SRSF1 | GGAGGRV | 4572 | 18.04 | 4380 | 17.29 | 1.04 | 7.3 | 2.18e-2 | 2.78e0 | 3.08e-2 |
| All targets | RNCMPT00106 | SRSF1 | GGAGGAM | 3157 | 12.46 | 2998 | 11.83 | 1.05 | 8.3 | 2.20e-2 | 2.82e0 | 3.08e-2 |
| All targets | RNCMPT00108 | SRSF1 | GGAGGRV | 4046 | 15.97 | 3883 | 15.32 | 1.04 | 7.6 | 3.44e-2 | 4.41e0 | 4.72e-2 |
| All targets | RNCMPT00116 | YBX1 | AACAUCA | 648 | 2.56 | 584 | 2.30 | 1.11 | 11 | 3.63e-2 | 4.65e0 | 4.88e-2 |
| D0-unique | RNCMPT00160 | HNRNPH2 | GGGAGGG | 1007 | 9.35 | 719 | 6.68 | 1.4 | 8.6 | 2.20e-12 | 2.81e-10 | 1.76e-10 |
| D0-unique | RNCMPT00088 | SRSF10 | AGAGARR | 45 | 0.42 | 4 | 0.04 | 9.2 | 10 | 4.11e-10 | 5.26e-8 | 1.10e-8 |
| D0-unique | RNCMPT00089 | SRSF10 | AGAGARR | 45 | 0.42 | 4 | 0.04 | 9.2 | 10 | 4.11e-10 | 5.26e-8 | 1.10e-8 |
| D0-unique | RNCMPT00067 | SRSF9 | GGRWGGA | 700 | 6.50 | 504 | 4.68 | 1.39 | 8.9 | 8.92e-9 | 1.14e-6 | 1.79e-7 |
| D0-unique | RBPMap-ESRP1-1 | ESRP1 | GGGKGG | 913 | 8.48 | 717 | 6.66 | 1.27 | 8.2 | 6.64e-7 | 8.50e-5 | 1.07e-5 |
| D0-unique | RBPMap-FUBP1-2 | FUBP1 | UDUUU | 1708 | 15.86 | 1447 | 13.44 | 1.18 | 7.5 | 1.82e-6 | 2.32e-4 | 2.43e-5 |
| D0-unique | RNCMPT00044 | PCBP2 | CCUYCCC | 256 | 2.38 | 166 | 1.54 | 1.54 | 9.5 | 6.87e-6 | 8.80e-4 | 7.50e-5 |

| | | | | | | | | | | | | |
|---|---|---|---|---|---|---|---|---|---|---|---|---|
| D0-unique | RNCMPT00117 | HuR | UUUGUUU | 579 | 5.38 | 440 | 4.09 | 1.32 | 9.2 | 7.48e-6 | 9.57e-4 | 7.50e-5 |
| D0-unique | RNCMPT00090 | SRSF10 | AGAGAMA | 139 | 1.29 | 76 | 0.71 | 1.82 | 10 | 1.04e-5 | 1.33e-3 | 9.24e-5 |
| D0-unique | RNCMPT00056 | RBM8A | GCGCGCG | 229 | 2.13 | 150 | 1.39 | 1.52 | 11 | 2.91e-5 | 3.72e-3 | 2.34e-4 |
| D0-unique | RNCMPT00052 | RBM4 | GCGCGGG | 150 | 1.39 | 88 | 0.82 | 1.7 | 13 | 3.52e-5 | 4.51e-3 | 2.43e-4 |
| D0-unique | RBPMap-EIF4G2-3 | EIF4G2 | GGUYGC | 639 | 5.94 | 504 | 4.68 | 1.27 | 10 | 3.63e-5 | 4.64e-3 | 2.43e-4 |
| D0-unique | RBPMap-BOLL-2 | BOLL | UUUDUUU | 1938 | 18.00 | 1704 | 15.83 | 1.14 | 6.2 | 5.62e-5 | 7.19e-3 | 3.47e-4 |
| D0-unique | RBPMap-EWSR1-2 | EWSR1 | GGGDGGGG | 1261 | 11.71 | 1081 | 10.04 | 1.17 | 7.6 | 1.08e-4 | 1.38e-2 | 6.17e-4 |
| D0-unique | RNCMPT00113 | RBM4 | GCGCGSG | 341 | 3.17 | 251 | 2.33 | 1.36 | 10 | 1.24e-4 | 1.59e-2 | 6.64e-4 |
| D0-unique | RNCMPT00112 | HuR | UUUGUUU | 495 | 4.60 | 389 | 3.61 | 1.27 | 9.6 | 2.04e-4 | 2.61e-2 | 1.02e-3 |
| D0-unique | RBPMap-FUS-1 | FUS | GGKGG | 246 | 2.28 | 174 | 1.62 | 1.41 | 10 | 2.58e-4 | 3.31e-2 | 1.22e-3 |
| D0-unique | RNCMPT00176 | MSI1 | UAGUWRG | 2851 | 26.48 | 2596 | 24.11 | 1.1 | 6.5 | 2.89e-4 | 3.69e-2 | 1.29e-3 |
| D0-unique | RNCMPT00032 | HuR | UUWUUUU | 879 | 8.16 | 742 | 6.89 | 1.18 | 8.8 | 3.63e-4 | 4.64e-2 | 1.53e-3 |
| D0-unique | RNCMPT00036 | LIN28A | HGGAGAA | 712 | 6.61 | 592 | 5.50 | 1.2 | 8.7 | 4.88e-4 | 6.24e-2 | 1.96e-3 |
| D0-unique | RNCMPT00070 | SNRNP70 | RWUCAAG | 244 | 2.27 | 177 | 1.64 | 1.38 | 11 | 6.35e-4 | 8.13e-2 | 2.38e-3 |
| D0-unique | RNCMPT00018 | FUS | UGCGCGC | 86 | 0.80 | 48 | 0.45 | 1.78 | 13 | 6.52e-4 | 8.34e-2 | 2.38e-3 |
| D0-unique | RNCMPT00041 | MSI1 | UAGUWRG | 2281 | 21.19 | 2071 | 19.24 | 1.1 | 7 | 7.66e-4 | 9.80e-2 | 2.67e-3 |
| D0-unique | RNCMPT00049 | RBM28 | GWGUAGW | 61 | 0.57 | 33 | 0.31 | 1.82 | 11 | 2.54e-3 | 3.25e-1 | 8.43e-3 |
| D0-unique | RNCMPT00026 | HNRNPK | CCAAMCC | 1050 | 9.75 | 925 | 8.59 | 1.13 | 7.8 | 2.63e-3 | 3.36e-1 | 8.43e-3 |
| D0-unique | RNCMPT00045 | PPRC1 | SSGCGCS | 458 | 4.25 | 379 | 3.52 | 1.21 | 9.5 | 3.49e-3 | 4.47e-1 | 1.08e-2 |
| D0-unique | RNCMPT00186 | PCBP1 | CCUWWCC | 1543 | 14.33 | 1397 | 12.98 | 1.1 | 7.5 | 3.74e-3 | 4.79e-1 | 1.10e-2 |
| D0-unique | RBPMap-CNOT4-1 | CNOT4 | ACACAG | 2420 | 22.48 | 2237 | 20.78 | 1.08 | 7.1 | 3.82e-3 | 4.89e-1 | 1.10e-2 |

| | | | | | | | | | | | | |
|---|---|---|---|---|---|---|---|---|---|---|---|---|
| D0-unique | RNCMPT00020 | FXR2 | GGACRRG | 3040 | 28.24 | 2838 | 26.36 | 1.07 | 5.5 | 4.37e-3 | 5.60e-1 | 1.21e-2 |
| D0-unique | RNCMPT00043 | PABPC4 | AAAAAAA | 402 | 3.73 | 332 | 3.08 | 1.21 | 8.8 | 5.41e-3 | 6.93e-1 | 1.45e-2 |
| D0-unique | RNCMPT00185 | KHDRBS2 | RAUAAAM | 2128 | 19.77 | 1967 | 18.27 | 1.08 | 7.1 | 6.20e-3 | 7.94e-1 | 1.61e-2 |
| D0-unique | RNCMPT00136 | HuR | UUGGUUU | 738 | 6.85 | 645 | 5.99 | 1.14 | 9 | 6.67e-3 | 8.54e-1 | 1.63e-2 |
| D0-unique | RNCMPT00110 | SRSF1 | MAGGACAV | 1673 | 15.54 | 1532 | 14.23 | 1.09 | 7.5 | 6.69e-3 | 8.57e-1 | 1.63e-2 |
| D0-unique | RNCMPT00086 | ZC3H14 | UUUGUUU | 1812 | 16.83 | 1668 | 15.49 | 1.09 | 7.3 | 7.67e-3 | 9.81e-1 | 1.79e-2 |
| D0-unique | RNCMPT00073 | SRSF7 | GACGACGR | 328 | 3.05 | 268 | 2.49 | 1.22 | 8.9 | 7.79e-3 | 9.98e-1 | 1.79e-2 |
| D0-unique | RNCMPT00050 | RBM3 | GAUACGA | 1790 | 16.63 | 1649 | 15.32 | 1.09 | 6.8 | 8.48e-3 | 1.09e0 | 1.89e-2 |
| D0-unique | RNCMPT00154 | RBM5 | GAAGGAG | 461 | 4.28 | 394 | 3.66 | 1.17 | 9.4 | 1.20e-2 | 1.53e0 | 2.60e-2 |
| D0-unique | RNCMPT00063 | SAMD4A | GCUGGMC | 1093 | 10.15 | 990 | 9.20 | 1.1 | 8.3 | 1.27e-2 | 1.63e0 | 2.68e-2 |
| D0-unique | RNCMPT00083 | YBX1 | AACAUCA | 1481 | 13.76 | 1365 | 12.68 | 1.08 | 6.8 | 1.55e-2 | 1.99e0 | 3.20e-2 |
| D0-unique | RNCMPT00172 | IGF2BP3 | AMAHWCA | 958 | 8.90 | 872 | 8.10 | 1.1 | 7.5 | 2.34e-2 | 3.00e0 | 4.70e-2 |
| Shared targets | RBPMap-FUBP1-2 | FUBP1 | UDUUU | 3364 | 31.90 | 2819 | 26.74 | 1.19 | 7 | 2.22e-12 | 2.84e-10 | 1.89e-10 |
| Shared targets | RNCMPT00032 | HuR | UUWUUUU | 867 | 8.22 | 649 | 6.16 | 1.34 | 9.1 | 1.19e-8 | 1.52e-6 | 5.06e-7 |
| Shared targets | RNCMPT00160 | HNRNPH2 | GGGAGGG | 1736 | 16.46 | 1427 | 13.53 | 1.22 | 7.9 | 2.12e-8 | 2.71e-6 | 6.03e-7 |
| Shared targets | RNCMPT00044 | PCBP2 | CCUYCCC | 360 | 3.41 | 230 | 2.18 | 1.56 | 9.4 | 4.87e-8 | 6.23e-6 | 1.04e-6 |

| | | | | | | | | | | | | |
|---|---|---|---|---|---|---|---|---|---|---|---|---|
| Shared targets | RNCMPT00086 | ZC3H14 | UUUGUUU | 1091 | 10.35 | 873 | 8.28 | 1.25 | 8.1 | 4.76e-7 | 6.09e-5 | 6.99e-6 |
| Shared targets | RNCMPT00150 | ESRP2 | UGGGGAU | 650 | 6.16 | 485 | 4.60 | 1.34 | 9 | 5.41e-7 | 6.92e-5 | 6.99e-6 |
| Shared targets | RNCMPT00156 | CNOT4 | GACAGAN | 63 | 0.60 | 19 | 0.18 | 3.2 | 12 | 5.73e-7 | 7.34e-5 | 6.99e-6 |
| Shared targets | RNCMPT00117 | HuR | UUUGUUU | 347 | 3.29 | 234 | 2.22 | 1.48 | 9.5 | 1.58e-6 | 2.02e-4 | 1.68e-5 |
| Shared targets | RBPMap-BOLL-2 | BOLL | UUUDUUU | 754 | 7.15 | 589 | 5.59 | 1.28 | 6.9 | 3.72e-6 | 4.77e-4 | 3.53e-5 |
| Shared targets | RNCMPT00112 | HuR | UUUGUUU | 1745 | 16.55 | 1522 | 14.43 | 1.15 | 7.1 | 5.11e-5 | 6.54e-3 | 4.04e-4 |
| Shared targets | RNCMPT00049 | RBM28 | GWGUAGW | 78 | 0.74 | 36 | 0.34 | 2.14 | 12 | 5.21e-5 | 6.67e-3 | 4.04e-4 |
| Shared targets | RNCMPT00070 | SNRNP70 | RWUCAAG | 625 | 5.93 | 501 | 4.75 | 1.25 | 9.8 | 1.22e-4 | 1.56e-2 | 8.04e-4 |
| Shared targets | RNCMPT00176 | MSI1 | UAGUWRG | 2154 | 20.43 | 1919 | 18.20 | 1.12 | 7.3 | 1.22e-4 | 1.57e-2 | 8.04e-4 |
| Shared targets | RBPMap-ELAVL4-1 | ELAVL4 | UAAUU | 1161 | 11.01 | 998 | 9.47 | 1.16 | 9.5 | 2.43e-4 | 3.11e-2 | 1.48e-3 |
| Shared targets | RBPMap-FUS-1 | FUS | GGKGG | 141 | 1.34 | 94 | 0.89 | 1.49 | 11 | 1.31e-3 | 1.68e-1 | 7.34e-3 |

| | | | | | | | | | | | | |
|---|---|---|---|---|---|---|---|---|---|---|---|---|
| Shared targets | RNCMPT00172 | IGF2BP3 | AMAHWCA | 1898 | 18.00 | 1717 | 16.28 | 1.11 | 6.9 | 1.38e-3 | 1.76e-1 | 7.34e-3 |
| Shared targets | RNCMPT00084 | YBX2 | AACAWCD | 877 | 8.32 | 756 | 7.17 | 1.16 | 9.2 | 1.49e-3 | 1.90e-1 | 7.46e-3 |
| Shared targets | RNCMPT00108 | SRSF1 | GGAGGRV | 2199 | 20.86 | 2016 | 19.12 | 1.09 | 7.1 | 2.53e-3 | 3.23e-1 | 1.20e-2 |
| Shared targets | RNCMPT00045 | PPRC1 | SSGCGCS | 352 | 3.34 | 284 | 2.69 | 1.24 | 10 | 3.92e-3 | 5.02e-1 | 1.76e-2 |
| Shared targets | RNCMPT00109 | SRSF1 | GGAGGRV | 2299 | 21.80 | 2128 | 20.18 | 1.08 | 6.9 | 5.31e-3 | 6.79e-1 | 2.09e-2 |
| Shared targets | RNCMPT00090 | SRSF10 | AGAGAMA | 235 | 2.23 | 182 | 1.73 | 1.29 | 10 | 5.40e-3 | 6.91e-1 | 2.09e-2 |
| Shared targets | RNCMPT00033 | IGF2BP2 | AMAWACA | 557 | 5.28 | 475 | 4.50 | 1.17 | 8.4 | 5.83e-3 | 7.46e-1 | 2.09e-2 |
| Shared targets | RNCMPT00021 | G3BP2 | AGGAURA | 346 | 3.28 | 282 | 2.67 | 1.23 | 10 | 5.94e-3 | 7.60e-1 | 2.09e-2 |
| Shared targets | RNCMPT00163 | SRSF1 | GGAGGAG | 1987 | 18.84 | 1831 | 17.37 | 1.09 | 6.9 | 6.06e-3 | 7.75e-1 | 2.09e-2 |
| Shared targets | RNCMPT00041 | MSI1 | UAGUWRG | 2673 | 25.35 | 2492 | 23.63 | 1.07 | 6.8 | 6.13e-3 | 7.84e-1 | 2.09e-2 |
| Shared targets | RNCMPT00107 | SRSF1 | GGAGGAN | 3869 | 36.69 | 3654 | 34.65 | 1.06 | 4.8 | 6.80e-3 | 8.71e-1 | 2.23e-2 |

| | | | | | | | | | | | | |
|---|---|---|---|---|---|---|---|---|---|---|---|---|
| Shared targets | RNCMPT00106 | SRSF1 | GGAGGAM | 4604 | 43.66 | 4385 | 41.59 | 1.05 | 4.8 | 1.07e-2 | 1.37e0 | 3.40e-2 |
| Shared targets | RNCMPT00055 | RBM5 | GAAGGAA | 840 | 7.97 | 748 | 7.09 | 1.12 | 8.6 | 1.12e-2 | 1.43e0 | 3.41e-2 |
| Shared targets | RNCMPT00113 | RBM4 | GCGCGSG | 526 | 4.99 | 454 | 4.31 | 1.16 | 8.9 | 1.16e-2 | 1.49e0 | 3.43e-2 |
| Shared targets | RNCMPT00056 | RBM8A | GCGCGCG | 221 | 2.10 | 176 | 1.67 | 1.25 | 11 | 1.35e-2 | 1.73e0 | 3.85e-2 |
| Shared targets | RBPMap-ESRP1-1 | ESRP1 | GGGKGG | 204 | 1.93 | 162 | 1.54 | 1.26 | 9.6 | 1.60e-2 | 2.05e0 | 4.30e-2 |
| Shared targets | RNCMPT00067 | SRSF9 | GGRWGGA | 3825 | 36.28 | 3639 | 34.51 | 1.05 | 6 | 1.61e-2 | 2.06e0 | 4.30e-2 |

**Table S11.** Differential gene enrichment results from DESeq2 analysis of fRIP-Seq data at D0 (Sheet 1) and D28 (Sheet 2). See separate file.

**Table S12.** Integration of differential methylation results from DSS analysis of WGBS data at D0 and D7 with fRIP-Seq, IP-MS, CUT&RUN and RNA-Seq datasets. Sheet 1: Differentially methylated regions (DMRs) and their genomic annotation at D0. Sheet 2: DMRs and their genomic annotation at D7. Sheet 3: Promoter associated DMRs at D0 and D7 that overlap H3K4me/H3K27me3 CUT&RUN peaks ('CUT&RUN_Overlap' column), encode RNAs that are HNRNPU targets or proteins interacting with HNRNPU ('DMR_PPI_fRIP_Overlap' column), or show differential RNA abundance at D28 after HNRNPU silencing based on our previous bulk RNA-Seq data. The 'DMR_DEG_Overlap' column indicates whether the associated RNA is up- or downregulated, and the 'DEG_Concordant' column denotes whether the direction of RNA abundance change is concordant with the methylation change (TRUE/FALSE). See separate file.
